# Single-cell repertoire tracing identifies rituximab refractory B cells during myasthenia gravis relapses

**DOI:** 10.1101/840389

**Authors:** Ruoyi Jiang, Miriam L. Fichtner, Kenneth B. Hoehn, Panos Stathopoulos, Richard J. Nowak, Steven H. Kleinstein, Kevin C. O’Connor

## Abstract

Rituximab, a B cell-depleting therapy, is indicated for treating a growing number of autoantibody-mediated autoimmune disorders. However, relapses can occur after treatment and autoantibody-producing B cell subsets may be found during relapses. It is not understood if these autoantibody-producing B cell subsets emerge from the failed depletion of pre-existing B cells or are re-generated *de novo*. To further define the mechanisms that cause post-rituximab relapse, we studied patients with autoantibody-mediated muscle-specific kinase (MuSK) myasthenia gravis (MG) who relapsed after treatment. We carried out single-cell transcriptional and B cell receptor (BCR) profiling on longitudinal B cell samples. We identified clones present prior to therapy that continued to persist during relapse. Persistent B cell clones included both antibody-secreting cells and memory B cells characterized by gene expression signatures associated with B cell survival. A subset of persistent antibody-secreting cells and memory B cells were specific for the MuSK autoantigen. These results demonstrate that rituximab is not fully effective at eliminating autoantibody-producing B cells and provide a mechanistic understanding of post-rituximab relapse in MuSK MG.

## Introduction

Acquired myasthenia gravis (MG) is an autoimmune disorder caused by pathogenic autoantibodies that bind to membrane proteins at the neuromuscular junction leading to muscle weakness (1). While the majority of MG patients have serum autoantibodies against the acetylcholine receptor (AChR), a subset of patients have autoantibodies against muscle-specific tyrosine kinase (MuSK) (2, 3). MuSK is essential for AChR clustering and synaptic differentiation (4). Through binding MuSK, serum-derived autoantibodies impair clustering of AChRs, and block neuromuscular transmission causing disease (5).

Anti-CD20 mediated B cell depletion therapy (BCDT) is indicated for the treatment of multiple autoimmune disorders including rheumatoid arthritis (RA), autoimmune-associated vasculitides (AAV) and multiple sclerosis (MS) (6–8). The conspicuous clinical benefit of BCDT in patients with MuSK MG was recently demonstrated (9–11). However, a fraction of patients treated with the anti-CD20 BCDT agent rituximab (RTX) experienced post-rituximab (post-RTX) relapses despite symptomatic responses and a fall in MuSK autoantibody titer (9, 12–14).

Post-RTX relapse has been reported in the context of RA, AAV and systemic lupus erythematosus (SLE) (15–17). Relapses are associated with an elevated fraction of antigen-experienced B cells after depletion and are generally managed with additional BCDT (18–23). Post-RTX relapse in MuSK MG is associated with the presence of circulating plasmablast expansions, which include those that secrete MuSK-specific autoantibodies (24, 25). Previous studies have shown that antigen-experienced plasma cells or plasmablasts are resistant to BCDT owing to their low CD20 expression (26–29). Moreover, memory B cells, which express CD20, are also abundant after BCDT, and their increased frequency is associated with post-RTX relapses (19, 20). Whether or not plasmablasts, plasma cells or memory B cells emerge from inadequately depleted circulating B cells (versus emerging entirely *de novo*) is not known. The characteristics of these antigen-experienced B cells that resist depletion also remain unclear.

Understanding the mechanisms underlying post-RTX relapse is important for the development of new modalities for monitoring depletion and more effective therapy combinations for treating MG and other autoimmune disorders. Accordingly, we sought to further understand whether post-RTX relapse in MuSK MG was associated with the failed depletion of pre-existing B cells, including autoantibody-producing B cell subsets. In addition, we investigated the phenotype of the B cells that persist throughout the course treatment and relapse. To this end, we integrated data from multiple profiling methods on the same patients, including: (1) adaptive immune receptor repertoire sequencing (AIRR-Seq) to generate high-depth BCR libraries from unselected PBMCs collected at pre-RTX treatment time points and during episodes of post-RTX relapse; (2) previously published sequences of human monoclonal autoantibodies, with known specificity for the MuSK autoantigen, that emerged during post-RTX relapses; and (3) single-cell gene expression analysis with paired BCR repertoire sequencing, to trace B cell populations temporally and to identify the phenotype of B cells that emerge during post-RTX relapse. In comparison to the examination of clonal overlap between bulk BCR repertoires alone, this approach, which we refer to as Single-cell Tracing of Adaptive Immune Receptor repertoires (STAIR), allows for the unbiased characterization of the gene expression profile of persistent, disease-relevant B cells (including those specific for the autoantigen) at single-cell resolution. By using this approach, we offer new insights into the mechanism of post-RTX relapse in patients with MuSK MG, and potential avenues for improving the treatment of MG and other autoantibody-driven autoimmune diseases.

## Results

### Experimental design

We used the following experimental approach (Figure 1) to identify and characterize B cell clones that resist RTX depletion in three MuSK MG patients who experienced relapses. We first tested whether B cells clones persisted across pre- and post-RTX repertoires. To accomplish this, we sequenced the BCR repertoire of unselected PBMCs at high-depth using a bulk approach to capture a large number of sequences for identifying persistent clones. Second, we tested whether MuSK autoantibody producing B cells were among these persistent clones. In a previous study of these same patients, we produced recombinant human monoclonal antibodies (24, 25), which were specific for MuSK, from single B cells isolated during post-RTX relapses. Here, we sought to determine whether those specific BCR clones were present in pre-RTX repertoires. Third, we identified the phenotype of both persistent and non-persistent B cell clones. This was achieved through paired transcriptional profiling and BCR sequencing of single cells. These BCR sequences were traced to high-depth bulk sequencing repertoires derived from pre-RTX timepoints. The transcriptional profiles of persistent and non-persistent B cell clones were then compared.

**Figure 1.**
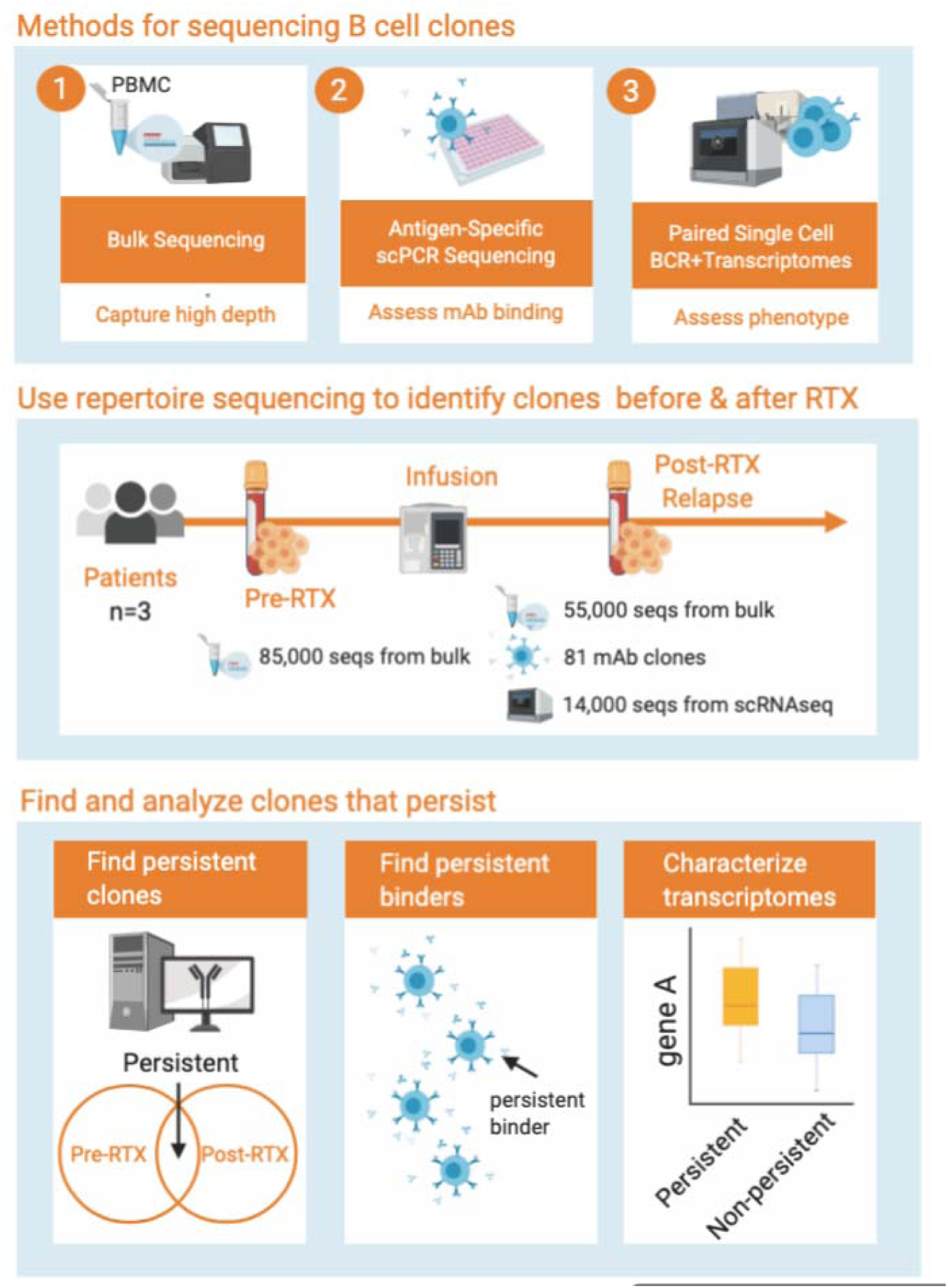
Schematic diagram showing overall workflow from clinical data elements and sample collection to computational analysis of BCR repertoires. Three approaches were used for sequencing. First, bulk repertoires using next-generation sequencing by isolating unpaired IGH V(D)J sequences directly from RNA were generated. Second, paired heavy and light chain V(D)J sequences from circulating B cells expressed as recombinant antibodies and tested for antigen specificity were traced to pre-RTX repertoires. Finally, paired transcriptome and heavy and light chain V(D)J sequence repertoires from high-throughput emulsion-based single cell sequencing was performed. These three types of repertoires were analyzed together in our analysis workflow for each patient across the sampled time points.

### A subset of B cell clones is refractory to depletion

We first sought to identify clones that escaped RTX depletion and contribute to relapses when the B cell compartment repopulates after treatment. To that end, we sequenced the B cell repertoire, using our high-depth bulk approach, from three MG patients before and after treatment (Table S1, S2). The number of V(D)J sequences isolated for patient 1 was 639,715; 82,720 from patient 2; and 107,504 from patient 3. Grouping the sequences into clones—those with a common ancestor based on their V, J gene usage and junction sequences (30)—generated 187,195; 34,719; and 14,579 clones for patient 1, 2 and 3 respectively. A total of 2,462 clones were found with members that were present in both the pre-rituximab (pre-RTX) and post-rituximab relapse (post-RTX) repertoire (termed persistent clones) in patient 1 (or 1.6% of pre-RTX clones); 902 (4.9%) and 345 (7.6%) were found in patients 2 and 3 respectively (Figure 2A). Given that clonal relationships are inferred computationally, we next sought to substantiate that these persistent clones were not the result of sequence similarity that happened by chance. Because clones cannot be shared across multiple individuals, the average number of clones shared between patients was calculated e.g. the average number of shared clones between patient 1 and 2, and patient 1 and 3 was calculated. Patient 1 shared an average of 0.4% clones in comparisons with other patients, patient 2 shared an average of 1.4%, and patient 3 shared an average of 0.9%. Quantification using a clonal overlap metric demonstrated that the observed overlaps between pre-RTX and post-RTX repertoires were significantly higher relative to this background (t-test P=0.037) (Figure 2B). These collective data demonstrate that circulating B cell clones persist despite RTX-mediated depletion of the repertoire.

**Figure 2.**
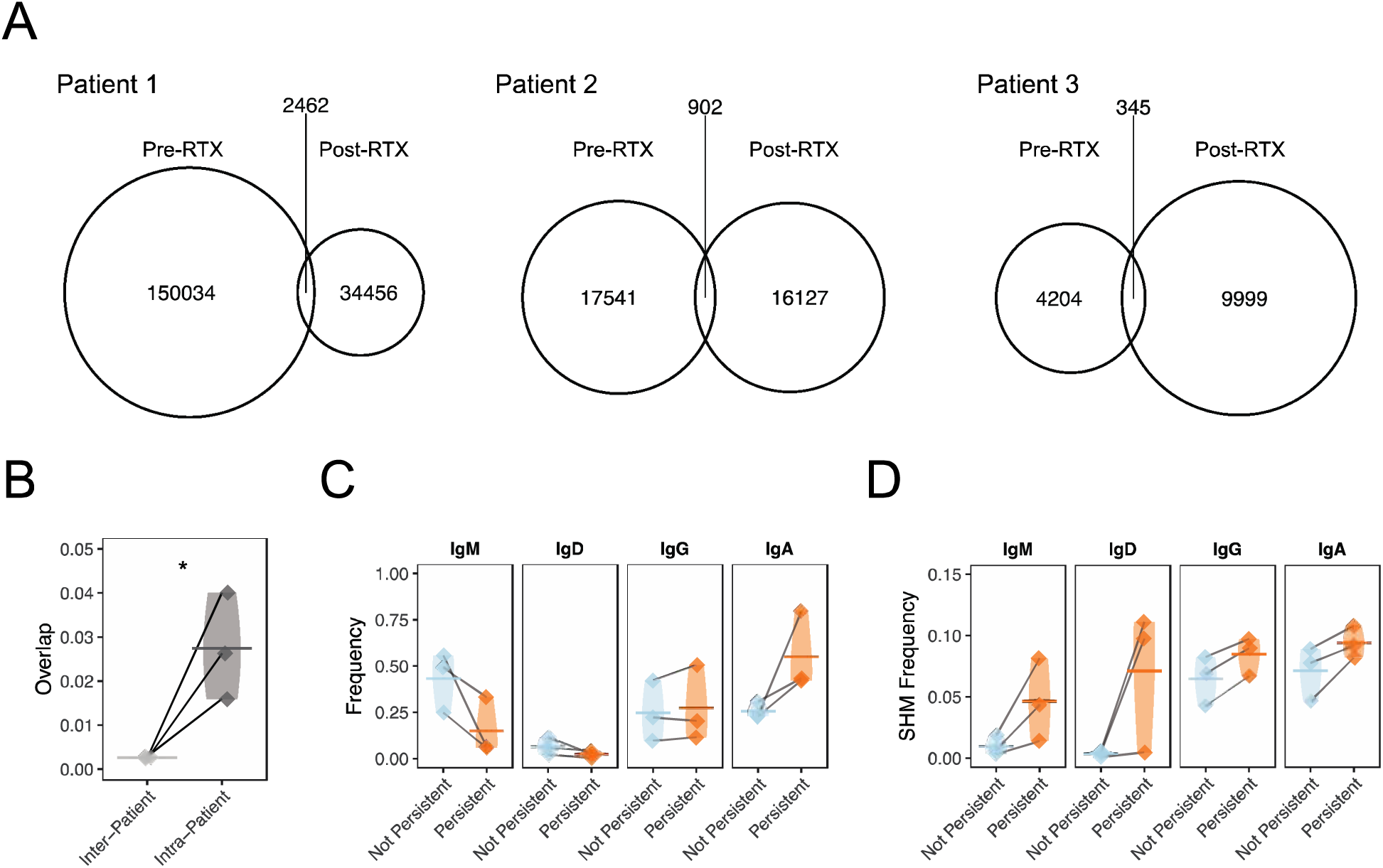
B cell clones that overlap pre-RTX and post-RTX bulk IGH repertoires (i.e., persistient clones) are associated with switched isoptypes and increased somatic hypermutation frequency. (A) Venn diagrams show counts of clones present at pre-RTX and post-RTX time points and those that overlap both time points for all study subjects (the relative area of each circle corresponds to the proportion of clones found at different time points for each patient). (B) Shared B cell clones between pre-RTX and post-RTX time points quantified as Bray-Curtis overlap for the same patient (intra-patient) or across different patients (inter-patient). Horizontal bars show the mean overlap of each comparison. A one-tailed t-test was used to assess the significance of the null hypothesis that intra-patient overlap was not higher than inter-patient overlap. Overall distribution of frequencies of different isotypes (C) and average somatic hypermutation frequencies (D) among the set of sequences belonging to persistent and non-persistent clones during post-RTX relapse for study subjects. Two-way ANOVA was performed to assess significance for an overall somatic hypermutation difference between non-persistent compared to persistent clones across isotype. Statistical differences are shown only when significant (****P < 0.0001; ***P < 0.001; **P < 0.01; *P<0.05).

### Persistent clones are antigen experienced

Next we compared the V(D)J properties of non-persistent clones to persistent clones that emerge post-RTX to investigate if the latter have distinct characteristics that might contribute to their persistence. Given the possible role of persistent B cells in recognizing antigen, we characterized features of the repertoire associated with antigen experience, namely isotype switching and elevated somatic hypermutation (SHM) frequency. We first examined our high-depth bulk repertoires; we quantified the difference in isotype frequency among V(D)J sequences from persistent and non-persistent post-RTX clones. V(D)J sequences from persistent clones were observed to be mostly isotype switched. While non-persistent sequences were 43.2% IgM, 6.4% IgD, 24.7% IgG and 25.5% IgA of the repertoire on average (Figure 2C), persistent ones were disproportionately switched (15.0% IgM, 2.2% IgD, 27.4% IgG and 55.0% IgA) (ANOVA p=0.027). In addition, the overall SHM frequency of persistent clones was significantly elevated (Figure 2D) (ANOVA P=0.014). We found that this difference in SHM frequency did not depend on the isotype of the sequence (ANOVA p = 0.287). Therefore, persistent clones are preferentially isotype switched with elevated SHM, reflecting a higher frequency of antigen-experienced clones.

### Persistent clones do not consistently expand or gain SHM

We next sought to determine whether antigen experience was gained during reconstitution of the B cell compartment after RTX. Examining the high-depth bulk repertiore, we first tested if persistent clones expand between pre-RTX and post-RTX time points. Clonal expansion was assessed by computing the change in size of persistent clones between pre-RTX and post-RTX time points. Average decreases of −11% (95% CI −93% to +1313%), −85% (95% CI −98% to +439%) and −7% (95% CI −99% to +40.5%) in clonal size respectively were observed across patients 1, 2 and 3 (one-sample t-test P=0.37) (Figure S3A-C). Thus, decreases in clonal size were observed but these decreases were not significant. Given that persistent clones have elevated frequencies of SHM, we tested whether individual clones acquire additional SHM by quantifying the average change in SHM frequency of persistent clones between pre-RTX and post-RTX time points. The median change in SHM frequency was −0.2% (95% CI −89% to +1132%), −2.4% (95% CI −74% to 187%) and +4.5% (95% CI −41% to +67%) for patients 1, 2 and 3 respectively (Figure S3D-F). Thus, we did not detect a consistent change in the SHM frequency of persistent clones between pre-RTX and post-RTX samples for these patients (one-sample t-test P=0.807). These collective findings suggest that clonal expansion and the acquisition of antigen experience between pre-RTX and post-RTX time points are not general features of persistent clones.

### Some persistent clones are disease relevant and bind MuSK

We then sought to investigate if persistent B cell clones were autoantigen-specific. In two previous studies, using the same three patients described here, we identified B cell clones during post-RTX relapse that were specific for autoantigen (Table S3). In the first study, 43 unique B cell clones were used to generate recombinant monoclonal antibodies, all of which were derived from circulating plasmablasts as this cell type was demonstrated to produce MuSK-specific antibodies when cultured *in vitro* (24). In the second, a fluorescent MuSK antigen bait was used to both identify and isolate MuSK-specific B cell clones from which an additional 38 recombinant monoclonal antibodies were produced (25). In total, 11 members of these collective 81 clones bound surface expressed MuSK across a range of different binding strengths when tested as monoclonal antibodies with a cell-based assay (24, 25). The BCR sequences of these clones were compared to the sequences in high-depth bulk repertiores to identify clonal relatives present both before and after treatment. Of these 81 clones, 27 were members of clones found in our bulk post-RTX BCR repertoires. This finding demonstrates that these clones were expanded, as they were found in unique, independent samples. When we tested if members of these 81 clones were present in pre-RTX repertoires, members belonging to 74 clones were not observed. However, we observed that seven clones could be found in pre-RTX repertoires. One of which was among the 11 demonstrated to specifically bind MuSK (Figure 3, Figure S4A-F) (24). Thus, disease-relevant and MuSK autoantigen-specific B cells present during relapse can derive from the failed depletion of B cell clones that existed prior to therapy.

**Figure 3.**
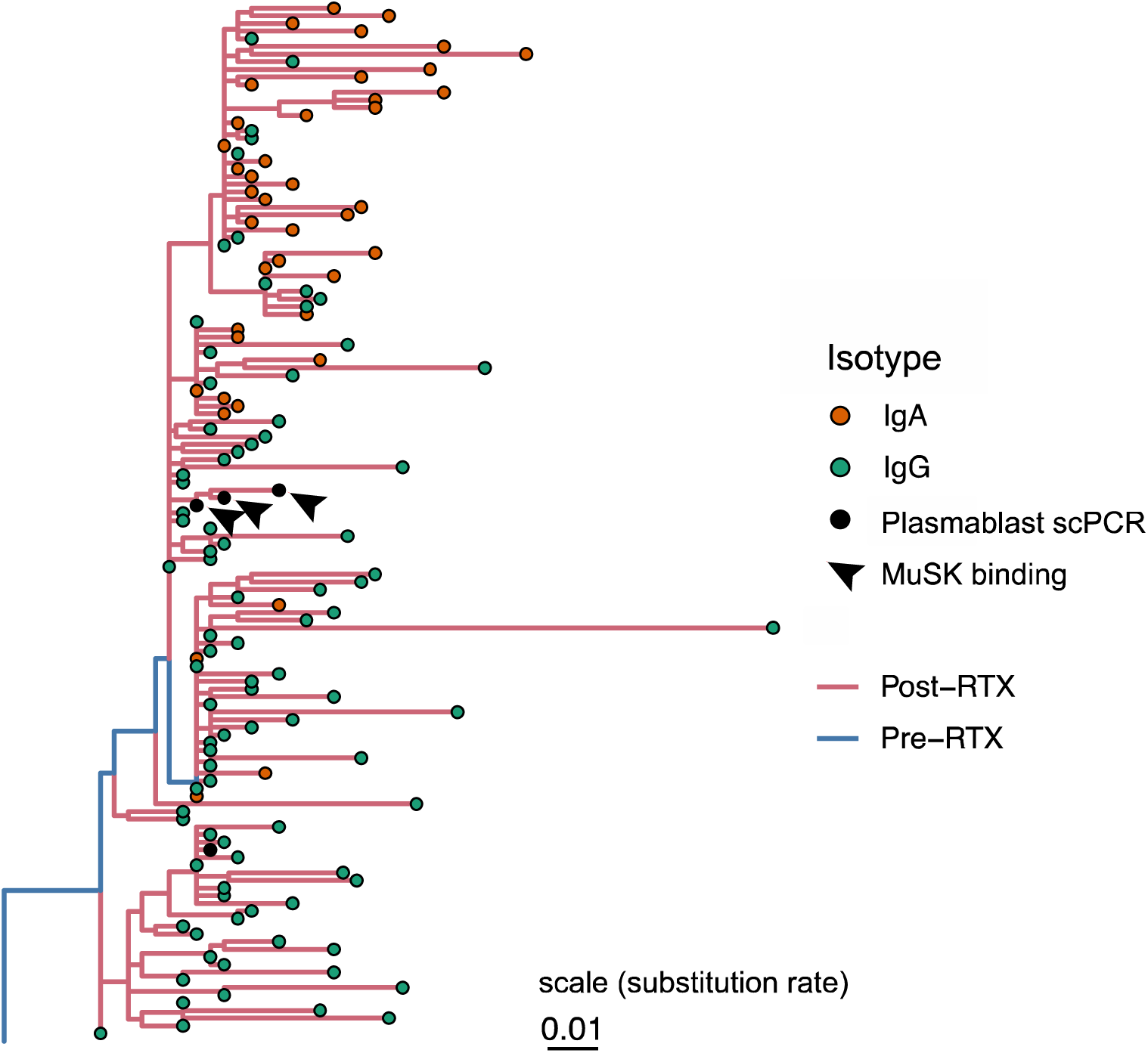
Example lineage tree of a disease-relevant B cell clone spanning both the pre- and post-RTX repertoire. Sequences isolated from plasmablasts using single-cell PCR (scPCR) sequencing-based approaches and tested as monoclonal antibodies are denoted as “Plasmablast scPCR” (black dots); three of these bind MuSK and are denoted as “MuSK binding” (black arrows). Lineage tree topology and branch lengths were estimated using maximum parsimony, with edge lengths representing the expected number of nucleotide substitutions per site (see scale) as estimated using dnapars v3.967 (88). Tips are colored by antibody isotype, and each internal branch is colored by whether its descendant node was determined to have occurred in the pre- or post-RTX repertoire using a constrained maximum parsimony algorithm (see Methods).

### Antibody-secreting cell-like B cells are preferentially persistent but memory B cells also persist

Given the disease relevance of persistent antigen-experienced B cells during relapse, we next sought to characterize the transcriptional state of persistent B cells to determine their phenotype in an unbiased manner. We also sought to quantify how different B cell subsets are represented among persistent populations. To accomplish this, we carried out single-cell transcriptome and paired BCR repertoire profiling of circulating post-RTX B cells. Owing to the high frequency of transitional B cells during post-RTX reconstitution, IgD^low^ B cells were selected through sorting prior to single-cell analysis (Figure S5). Further, the analysis was limited to the switched IgG and IgA repertoire by computationally filtering out cells with V(D)J’s paired with an IgD or IgM constant region. Clones were then identified in the single-cell data sets that were related to clones present in pre-RTX high-depth bulk BCR repertoires. This allowed for direct interrogation of the transcriptomes of persistent B cells. That is, we leveraged the resolution offered by high-throughput single-cell transcriptomes and high-depth offered by bulk BCR analysis to perform single-cell tracing of adaptive immune repertoires (STAIR). From single-cell sequencing, a total of 18,133 cells associated with 14,671 IGH V(D)J sequences were identified (Table S4) from the same three post-RTX patients and an additional asymptomatic AChR MG patient used as a control. Gene expression patterns were used to partition the cells into 11 distinct clusters (Figure S6A-E, Table S5). Clusters of antibody-secreting cell-like (ASC), memory B cells, mature naive B cells and transitional B cells were assigned by analysis of the gene expression of these clusters (Figure 4A). Because B cells from the single-cell expression analysis could not be experimentally isolated to assess for antibody secretion or other functional features associated with plasmablasts/plasma cells, we chose to label the cluster most associated with a plasmablast/plasma cell signature as antibody-secreting cell-like (hereafter referred to as ASC). By combining single-cell repertoires with high-depth bulk pre-RTX BCR repertoires, a total of 820 persistent clones with members in the single-cell profiling data were identified. Persistent clones could be identified in both the ASC as well as memory B cell clusters. The ASC cluster had a higher frequency (27.9%) of persistent clones in comparison to the memory cluster (14.6%) (Figure 4B, 4C) (t-test P=0.045). As further confirmation that some ASC and memory B cell clones are persistent, IGH sequences were also identified from single-cells in the ASC and memory B cell cluster with identical V(D)J sequence as pre-RTX members (Figure 4B, red dots). Thus, these data suggest that while both ASCs and memory B cells are persistent, members of the ASC subset are more likely to be found pre-RTX than memory B cells during relapse.

**Figure 4.**
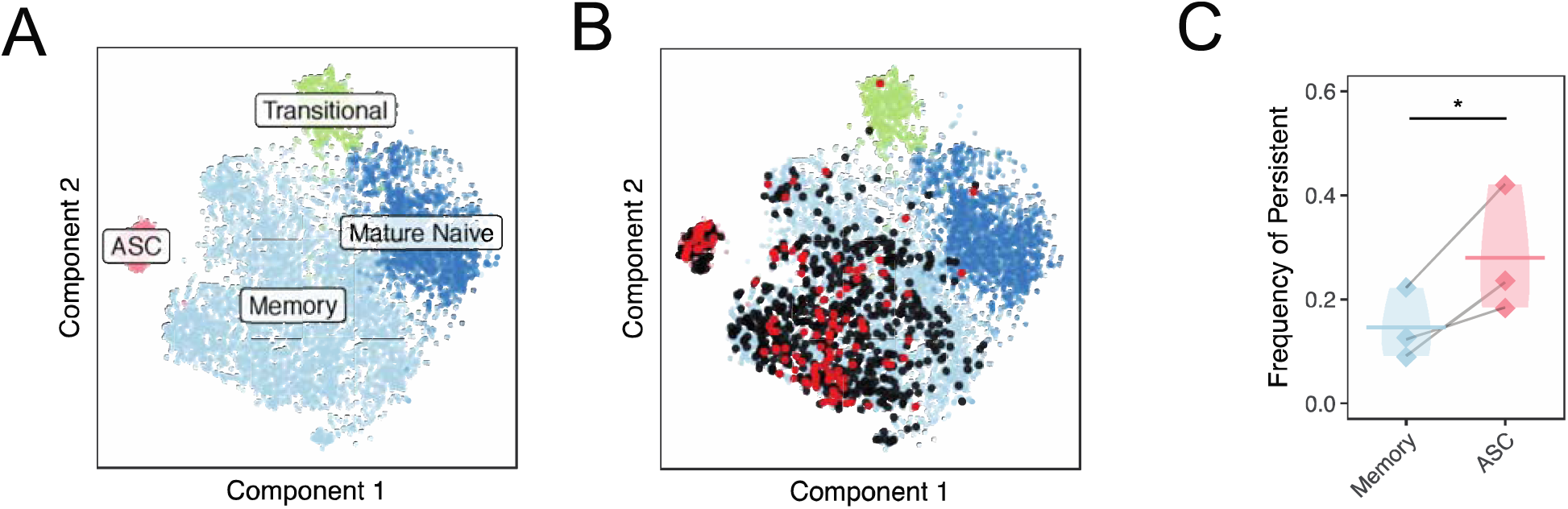
Persistent B cell clones are disproportionately represented by the ASC phenotype. (A) Unbiased clustering of single-cell transcriptomes from all study subjects is visualized by t-SNE and colored based on B cell subset assignment. (B) V(D)J sequences associated with persistent B cell clones are visualized over the tSNE plot labelled in (A). B cells are colored black or red if the heavy chain V(D)J sequence paired with the cell is part of a persistent B cell clone. B cells are colored red if the heavy chain V(D)J sequence paired with the cell has the exact same V(D)J sequence as one present in a pre-RTX bulk repertoire, and black if the clonal relative differs by one or more SHM. (C) The relative fractions of persistent compared to non-persistent B cells for the memory B cell cluster compared to ASC cluster. A one-tailed t-test was used to assess the significance of the null hypothesis that ASCs would not be enriched compared to memory B cells. Horizontal bars show the average frequency of a given B cell subset across patients. Frequencies belonging to the same patient are paired with a gray line. Statistical differences are shown only when significant (****P < 0.0001; ***P < 0.001; **P < 0.01; *P<0.05).

### ASCs have persistence-associated features compared to memory B cells

Given their role in the production of soluble immunoglobulin, ASCs can be disease-relevant in MG and other autoantibody-mediated diseases. Moreover, we previously demonstrated their direct role in MuSK autoantibody production (24, 25). We next sought to investigate phenotypic characteristics of ASCs compared to memory B cells during post-RTX relapse to further understand their mechanism of persistence and identify possible subsets of persistent ASCs. We first investigated the expression of CD20, the target of rituximab. To test the association between the expression of CD20 and persistence of ASCs, CD20 expression was examined among B cell subsets. The ASC cluster was observed to express lower CD20 levels when compared to memory B cells (t-test P=0.021) (Figure 5A). However, when we examined the expression of CD20 of persistent compared to non-persistent ASCs, a decrease was observed but this was not significant (t-test P=0.15) (Figure 5B). BAFF (also known as B cell activating factor, B lymphocyte stimulator or BlyS) is a TNF superfamily member produced almost exclusively by myeloid cells (31, 32). BAFF (and another TNF superfamily ligand, APRIL) can bind a family of B cell specific receptors to promote B cell survival. Given that APRIL/BAFF receptor genes have been observed to play a role in mediating B cell survival after RTX depletion, we investigated the differential expression of APRIL/BAFF receptor genes (33–35). Persistent ASCs were not observed to differentially express APRIL/BAFF receptor genes (TACI, BAFF-R and BCMA) (P=0.085, 0.555, 0.288 for TACI, BAFF-R, BCMA respectively).

**Figure 5.**
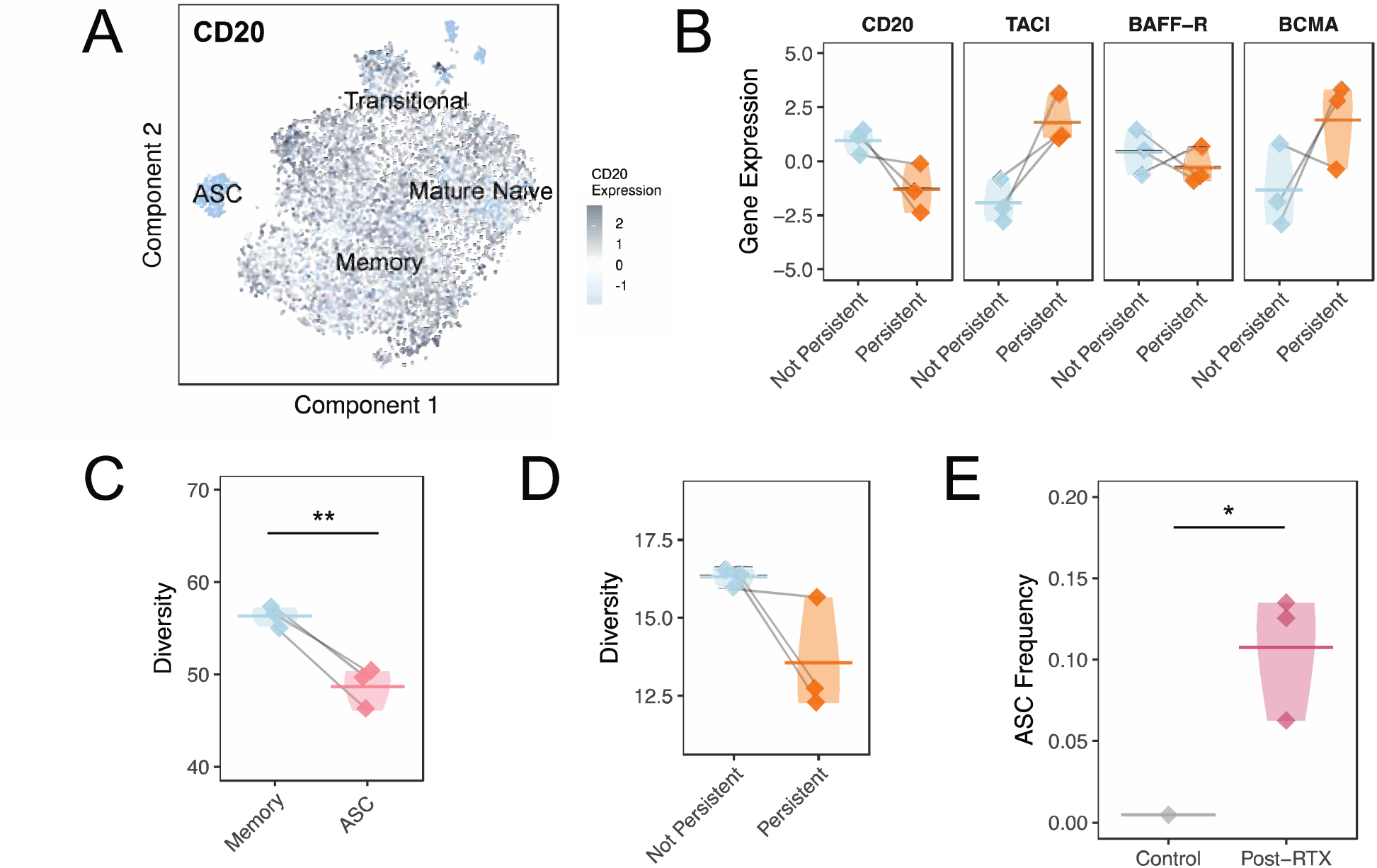
Unique characteristics of persistent ASCs. (A) Expression of log normalized CD20 expression visualized by intensity over t-SNE plot. (B) The expression of CD20 (the receptor target of rituximab gene symbol, MS4A1) and BAFF or APRIL receptors (TACI or TNFRSF13B; BAFF-R or TNFRSF13C; BCMA or TNFRSF17) for persistent and non-persistent members of the ASC cluster. Normalized gene expression values are computed from counts of gene expression transcripts. Paired two-tailed t-tests were used to test for the significant differential expression of each gene. Horizontal bars show the average expression of ASC cluster members for each patient for cells of a given status. (C) Clonal expansion expressed as Simpson’s diversity for ASC and memory B cell clusters. A one-tailed t-test was used to assess the significance of the null hypothesis that ASCs would not be less diverse than memory B cell clones. Horizontal bars show the average diversity of cluster members for each patient of memory B cells or ASCs. (D) Simpson’s diversity for persistent and non-persistent members of the ASC cluster. A one-tailed t-test was used to assess the significance of the null hypothesis that persistent clones would not be less diverse than non-persistent clones. Horizontal bars show the average diversity of ASC cluster members for each patient for cells of a given status. (E) The frequency of ASCs in post-RTX MuSK MG patients and a control AChR MG patient. The fraction of ASC cluster members is quantified as a ratio of of the number of cells in each cluster divided by the number of total B cells for each patient. A one sample t-test was used to assess the significance that ASCs would not be more abundant in post-RTX samples compared to the asymptomatic AChR MG patient sample. Horizontal bars show the average frequency of ASC cluster members for each patient of a given status. Values belonging to the same patient are paired with a gray line for all graphs. Statistical differences are shown only when significant (****P < 0.0001; ***P < 0.001; **P < 0.01; *P<0.05).

To investigate other features that might distinguish persistent and non-persistent ASCs, we examined their relative usage of different isotypes and average SHM frequencies. Persistent ASCs were also not observed to differ in terms of isotype usage (ANOVA P=0.194) or frequency of SHM (ANOVA P=0.101) compared to their non-persistent counterparts (Figure S7A, B). The clonal diversity (more clonal expansions) of the ASC cluster was lower compared to memory B cells (t-test P=0.011) (Figure 5C). However, when we examined clonal diversity within the ASC cluster, a consistent difference between persistent compared to non-persistent ASCs was not observed (t-test P=0.079) (Figure 5D). The fraction of cells in the ASC cluster was also observed to be 10.0% of the total repertoire in the post-RTX cohort (Figure 5E). This was significantly elevated compared to the 0.5% observed in the control asymptomatic AChR MG sample (one-sample t-test P=0.044). However, when we examined if clonal expansion of persistent B cells between the pre-RTX and post-RTX time points could explain the abundance of ASC B cells that we observed, ASC B cell clones were not consistently found to expand between pre-RTX and post-RTX time points with approximately half expanding and the other half shrinking; the average change in clonal size was therefore not consistent across patients (t-test P=0.099) (Figure S8A).

To summarize, persistent ASC cluster members were not observed to differ from non-persistent members in terms of their expression of CD20 and receptors for BAFF/APRIL, clonal diversity, isotype usage or SHM frequency. However, ASC cluster members expressed lower CD20 and were more clonally expanded than memory B cells overall. Collectively these data identify features of the ASC cluster as a whole compared to memory B cells that may explain their higher frequency among persistent clones.

### Persistent memory B cell clones are expanded and express low levels of CD20

Given the high frequency of persistent memory B cells observed, we next sought to explore if persistent memory B cells had distinct features compared to non-persistent memory B cells. We tested CD20 expression, expression of APRIL/BAFF receptors, SHM, isotype usage and clonal diversity. Persistent memory B cells were observed to have significantly lower CD20 expression (t-test P=0.022) and higher expression of TACI (t-test P=0.037), but lower expression of BAFF-R (t-test P=0.037) (Figure 6A). Analysis of SHM and isotype usage revealed that persistent memory B cells also had higher frequencies of SHM across isotypes (ANOVA P=0.037) (Figure 6B); the fraction of persistent memory B cells expressing IgG1 (t-test P=0.022) and IgG3 (t-test P=0.025) (ANOVA P= 0.007) was less than that of their non-persistent counterparts (Figure 6C). We examined if persistent memory B cells were clonally expanded compared to their non-persistent counterparts. They were observed to have lower clonal diversity post-RTX suggesting that they were clonally expanded (t-test P=0.030) (Figure 6D). Initial clustering of cells based on single-cell gene expression identified six distinct sub-clusters of memory B cells. To better characterize the gene expression features of persistent memory B cells (Figure S9A, B), we examined the most abundant sub-cluster among persistent memory B cells. This cluster (cluster 3) represented 36.7% of all persistent memory B cells but only 25.0% of all memory B cells (not the most prevalent overall) (Figure S9C). Analysis of gene expression showed that cluster 3 was characterized by lower CD20 expression (t-test P=0.005), higher TACI expression (t-test P=0.008) and lower BAFF-R expression (t-test P=0.014) compared to other memory B cell sub-clusters (Figure S10A). This cluster also displayed a higher frequency of SHM across all isotypes (ANOVA P=1.46e-6) (Figure S10C) and demonstrated increased usage of IgA compared to IgG isotypes (ANOVA P=0.0103) (Figure S10D), but was not more clonally expanded compared to other memory B cells (t-test P=0.169) (Figure S10E).

**Figure 6.**
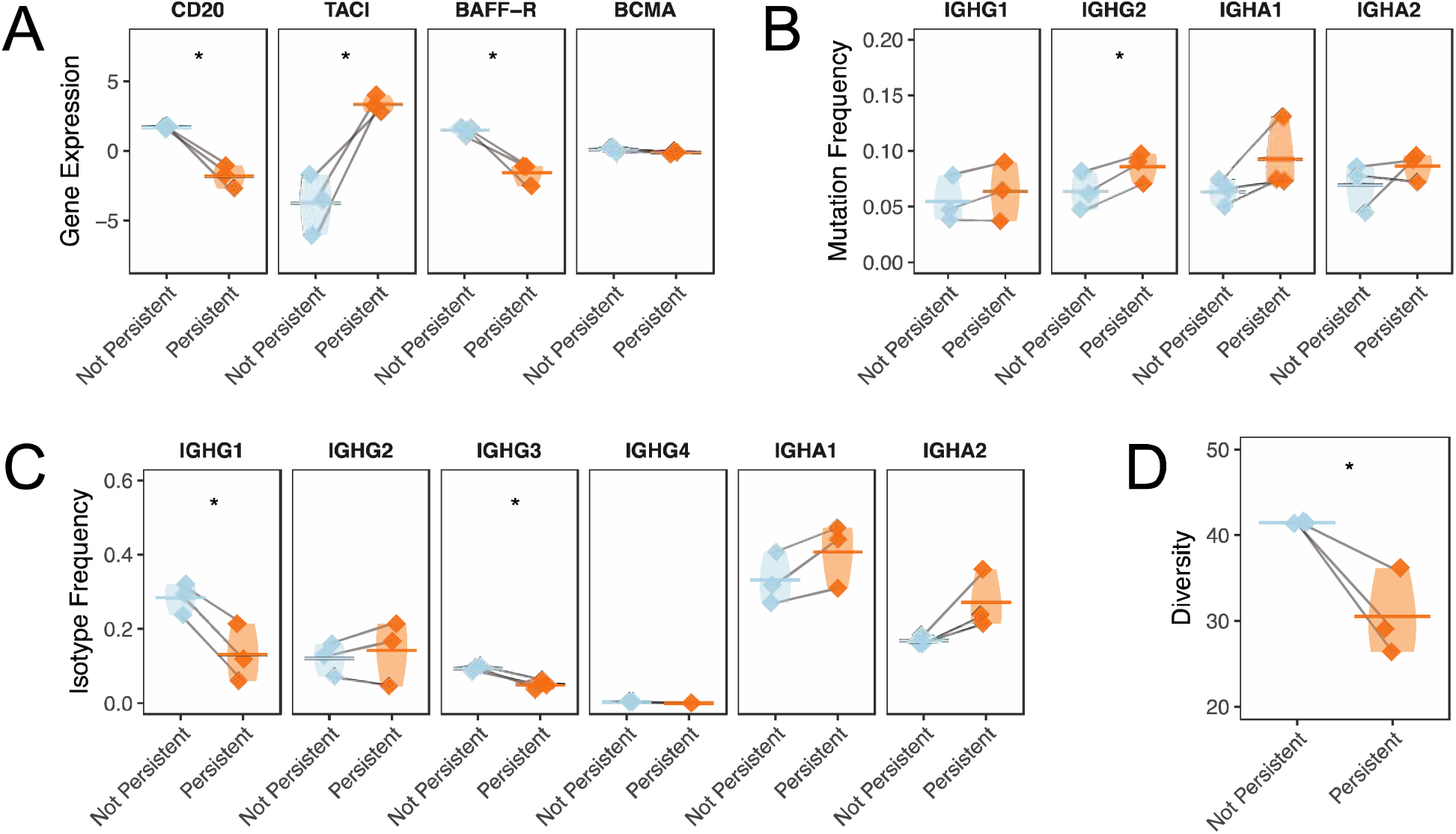
Memory B cells have distinct transcriptional and repertoire features associated with persistence. (A) The normalized gene expression of CD20 (the receptor target of rituximab gene symbol, MS4A1) and BAFF or APRIL receptors (TACI or TNFRSF13B; BAFF-R or TNFRSF13C; BCMA or TNFRSF17) is presented for persistent and non-persistent members of the memory B cell cluster for each patient for cells of a given status. Normalized gene expression values are computed from counts of gene expression transcripts. Paired two-tailed t-tests were used to test for the significant differential expression of each gene. (B) Individual SHM frequencies for persistent compared to non-persistent cells are presented for memory B cell cluster members. Only mean SHM frequencies are computed for isotypes with more than three V(D)J sequences. Horizontal bars show the average somatic hypermutation frequency for a given cluster. Paired two-tailed t-tests were used to assess the significance of differences in mutation frequency. (C) Overall constant region usage frequencies are quantified for persistent compared to non-persistent memory B cell cluster members per patient. Horizontal bars show the average frequency of constant region usage across patients. Two-way ANOVA was performed to assess significance for an overall isotype usage difference between non-persistent compared to persistent clones across isotype. Paired two-tailed t-tests were used to assess the significance for the differential usage of each constant region. (D) Clonal expansion expressed as Simpson’s diversity of persistent compared to non-persistent memory B cell cluster members. A one-tailed t-test was used to assess the significance of the null hypothesis that persistent clones would not be less diverse than non-persistent clones. Horizontal bars show the average diversity of memory B cell cluster members for each patient of a given status. Values belonging to the same patient are paired with a gray line for all graphs in this panel. Statistical differences are shown only when significant (****P < 0.0001; ***P < 0.001; **P < 0.01; *P<0.05).

Unbiased differential gene expression identified 1,282 differentially expressed genes between memory cluster 3 (enriched for persistent clones) and all other memory B cell sub-clusters (Data File S1) (FDR < 0.05). The upregulated subset of these genes was enriched for a MYC signature (or c-Myc, a transcription factor that enhances B cell survival) and BRCA1/p53 signature (transcription factors that regulate cell senescence and cell cycle) (Figure S11B) (36–38). The expression of two tissue homing genes that are the targets of the immunotherapy agents, natalizumab and fingolimod (ITGA4 and S1PR1 respectively), were also observed to be upregulated among cluster 3 memory B cells (P=0.049 for ITGA4 and P=0.039 for S1PR1) (Data File S1) (39). Thus, persistent memory B cells are clonally expanded and are characterized by a subset of B cells with elevated SHM, high expression of TACI, low expression of CD20 and a gene expression pattern associated with transcription factors that control cell survival, proliferation and senescence.

## Discussion

RTX is a B cell depleting agent widely used for the treatment of autoimmunity, however relapses after treatment have been reported in the context of diseases such as SLE, RA and AAVs (16, 17, 40, 41). MG is a chronic autoimmune disorder characterized by muscle weakness and caused by pathogenic autoantibodies that bind to membrane proteins at the neuromuscular junction (1). Pathogenic autoantibodies to MuSK can be found in patients with MG who do not have detectable antibodies to the acetylcholine receptor (2, 3). Recent studies have demonstrated the remarkable benefits of RTX-mediated B cell depletion therapy in MuSK MG (9, 10, 42). In addition to diminished autoantibody titers and significant clinical improvement, RTX also allows for a tapering and subsequent discontinuation of other immunotherapies in subsets of MG patients. However, post-RTX relapses do occur (9, 43). We recently demonstrated that relapses after RTX in MuSK MG patients are associated with the presence of circulating plasmablasts that express MuSK autoantibodies (24, 25). We sought to investigate if these disease-relevant and autoantigen-binding B cells derive either from the failed depletion of pre-existing or *de novo* B cell clones, and we further explored the characteristics of antigen-experienced B cells that resist B cell depletion in these patients.

We show that B cell clones, including MuSK-specific autoantibody-secreting B cells and others that may be disease-relevant, can emerge from the failed depletion of pre-existing B cell clones. Moreover, by directly tracing individual persistent B cell clones, we also show that persistent B cells more often use isotype-switched constant regions and have elevated frequencies of SHM. These are all features of an antigen-driven B cell response. We used a novel approach combining multiple methods of sequencing the BCR repertoire with single-cell gene expression (STAIR) to demonstrate that persistent B cell clones are comprised of both expanded ASCs as well as memory B cell populations expressing low levels of CD20 associated with a unique gene expression profile. Collectively, these data directly show that circulating ASC and memory B cells, including MuSK-specific B cells, are incompletely depleted by RTX in a subset of MG patients who experienced post-RTX relapses.

By using the BCR repertoire to directly trace and quantify the frequency of these ASCs, we show that many ASCs that reconstitute the B cell compartment after therapy come from pre-existing, circulating B cell populations detectable prior to depletion. This finding is consistent with a large body of evidence showing that ASCs resist depletion owing to their low levels of CD20 expression and tissue localization (26, 28, 29, 44). An elevated ASC frequency during post-RTX repopulation is also known to be a reliable predictor of poor treatment response in SLE and RA patients (15, 45–47). We also found that memory B cell clones can be identified that persist after RTX. The finding that circulating-memory B cell clones are also persistent is consistent with data showing that the frequency of memory populations after depletion predicts poor responses to RTX in SLE and RA (19–21). While these previous studies were important for identifying predictors of outcome, they were not designed to resolve B cell subsets due to limitations inherent with flow cytometry. By comparison, our definition of memory using single-cell transcriptomes does not rely on flow cytometry markers, but rather hundreds of markers that allow us to be confident these cells share common gene expression patterns associated with memory B cells. Furthermore, we were able to profile persistent memory B cell subsets in detail. For instance, a recent study using bulk AIRR-Seq to investigate the use of RTX for treating AAV, demonstrated that persistent clonotypes are isotype-switched, which we also observed in our study (48). However, because this analysis was not combined with single cell gene expression data, it is not clear if these results are confounded by the high frequency of certain B cell subsets that bear isotype-switched B cell receptors (48). In our study, persistence is correlated with a specific gene expression signature, that is, lower CD20 expression and differences in APRIL/BAFF receptor expression relative to other memory B cells.

Studies of RTX responses in anti-MAG-neuropathy (Immunoglobulin M(IgM) Anti-Myelin Associated Glycoprotein Peripheral Neuropathy) and pemphigus vulgaris (PV) (21, 23, 49) demonstrated that poor responses to RTX are associated with increased repertoire clonality post-RTX. Our results show that a large fraction of memory B cells are persistent, and that these persistent memory B cells are more clonally expanded than non-persistent memory B cells. These findings suggest that the association between increased repertoire clonality and poor responses to RTX may result from the persistence of clonally expanded memory B cells.

Our work is congruent with studies that have shown that BAFF and the differential expression of receptors for APRIL/BAFF (TACI, BAFF-R, BCMA) on antigen-experienced B cells may lead to poor outcomes after BCDT (33–35, 50). BAFF (also known as B cell activating factor, B lymphocyte stimulator or BlyS) is a TNF superfamily member produced almost exclusively by myeloid cells (31, 32). BAFF (and another TNF superfamily ligand, APRIL) can bind a family of B cell specific receptors to promote B cell survival. A higher frequency of memory B cells expressing lower BAFF-R can be detected in patients with RA relapsing after RTX (35) as well as in patients with GVHD who respond poorly to RTX (34). This is consistent with our observation that persistent memory B cells express lower BAFF-R levels. BAFF levels have been shown to negatively correlate with BAFF-R expression after RTX therapy in primary Sjogren’s syndrome and SLE (50). Moreover, extended exposure of B cells to soluble BAFF decreases surface levels of BAFF-R *in vitro*. By comparison, exposure to anti-BAFF agents has been shown to increase the expression of BAFF-R on B cells (51). By directly characterizing the features of B cell clones that resist RTX depletion we are able to show that memory B cell subsets that persist after RTX express surface markers that are associated with poor responses to RTX during relapse. These expression changes may thus reflect the underlying effectiveness of B cell depletion.

Our analysis of bulk BCR sequencing showed that the largest fraction of persistent B cells was switched to the IgA isotype. STAIR analysis showed that two subsets observed to be most persistent, ASCs and cluster 3 memory B cells were also predominantly IgA. We speculate that the abundance of IgA expressing B cells could be related to BAFF expression given the role of BAFF (and receptors like TACI) in inducing class switch recombination, particularly in the generation of gut resident IgA-switched B cells (52–54). The presence of IgA-switched circulating ASCs has been shown to be dependent on BAFF (55). Moreover, IgA secreting plasma cells were shown to reduce disease severity in an experimental autoimmune encephalomyelitis (EAE) mouse model, owing to their production of IL-10 (56). Thus, further characterization of how the induction of IgA producing B cell subsets by RTX may be involved in disease may be warranted as a means to investigate alternative mechanisms of therapeutic benefit from BCDT.

The conclusions of this study offer insights into possible therapeutic and diagnostic avenues for the management of patients with autoimmunity who experience relapses after RTX. Our findings regarding the BAFF/APRIL system suggest that combining RTX with BAFF pathway inhibitors (e.g. belimumab) could deplete populations that persist after RTX including those shown to produce autoantibodies. Combination strategies involving both belimumab and RTX have been successful in a phase 2a randomized clinical trial (RCT) in SLE which was followed with an ongoing phase 3 RCT (57, 58). Similar investigations may be warranted in MG.

We observed that up-regulated genes associated with the subset of memory B cells most resistant to depletion (cluster 3) included those involved in tissue homing (ITGA4, S1PR1). S1P receptor is a gene that is up-regulated in lymphocytes localized to lymph nodes and inflamed tissues and target of the immunotherapeutic, fingolimod. ITGA4 is a component of VLA-4, an integrin dimer also responsible for the localization of leukocytes to inflamed tissue and target of the biologic agent natalizumab (39). We speculate that this could reflect a possible tissue-associated origin for this particular B cell subset. This is consistent with evidence showing that memory B cells that resist RTX depletion can be identified in lymphnodes, spleen, bone marrow and tonsils (18, 19, 22, 59). These findings suggest that therapies such as by engineered T cell technology including chimeric-antigen receptor (CAR) or chimeric-autoantibody receptor (CAAR) T cell therapy (60, 61) and other methods for eliminating tissue-localized subsets may be well suited for preventing and treating relapses.

Existing NGS approaches for generating immune repertoires commonly begin with genomic DNA or mRNA and produce high-depth bulk repertoires of unpaired immunoglobulin or T cell receptor V(D)J sequences (62–64). More recently developed approaches use single-cell isolation techniques with high-throughput sequencing to construct repertoires, sometimes with paired gene expression (65–68). The use of these single cell-based approahes include studies on tumor-infiltrating lymphocytes from cancer biopsies, infiltrates from the brain of mice and B cells from the spleen after vaccination in mice (66–68). No such studies have investigated the circulating B cell repertoire in patients with autoimmune disease thusfar. Previous immune repertoire studies of BCDT have identified persistent expanded clones and associations between clonality and responder status, and single-cell transcriptome studies have recapitulated flow cytometry results showing increased frequencies of antigen-experienced B cells at the peak of depletion (69–72).

Our ability to extract biologically meaningful results from STAIR analysis suggests that tracing individual BCR sequences associated with disease relapse could provide a powerful tool for monitoring patient responses to immunotherapies that rely on the depletion of B cells and T cells. Moreover, in comparison to techniques that use bulk BCR repertoire sequencing to monitor therapy progress, this approach allows for the individual characterization of lymphocytes that resist therapy based on gene expression which may suggest alternative avenues for treatment (73).

Several limitations of this study should be considered. First, the single-cell gene expression of B cells was characterized only after RTX depletion. Determining the phenotype of B cells that resist RTX depletion prior to therapy may be of considerable interest as subsets prone to resisting B cell depletion prior to relapse could be identified and investigated. A recent study demonstrated that not only do B cell clones move between peripheral blood and cerebrospinal fluid over the course of immunotherapy in a subset of patients with MS, but many shift between memory and plasma cell characteristics (74). Similar studies using single-cell transcriptomics could allow for the identification of memory B cell subsets that give rise to autoantibody-producing plasmablasts during relapse. Second, our results rely on tracing B cell clones by analysis of their IGH V(D)J sequences alone. While we (and others) have demonstrated that clonal relationships can be reliably inferred given the diversity of IGH V(D)J sequences, false positives may occur (30, 75, 76). Third, owing to the abundance of transitional B cells that re-populate the repertoire during RTX relapse and our interest in studying antigen-experienced B cell subsets, we specifically excluded IgD+ expressing B cell subsets from our single-cell analysis. Nevertheless, a small fraction of B cell receptors paired with an IGHD constant region (3.2-11.4%) was found in our single-cell data. This is likely because flow-based sorting is not absolute in eliminating gated populations. Additionally, the expressed mRNA transcriptome may not always reflect the expression of surface markers used for sorting. Finally, although we previously generated a number of human recombinant MuSK-specific monoclonal antibodies from B cells present during post-RTX relapse, these monoclonal antibodies recognized MuSK with different binding capacities ranging from weak to very strong (24, 25). The one that we were successful in tracing to the pre-RTX repertoire, in the present study, was among those showing weaker binding capacity. Accordingly, we are only able to speculate whether it possessed pathogenic capacity *in vivo*. That it was a member of a large expanded clone with multiple clonal variants may suggest that this particular clonal member represented an intermediate step toward a higher binding capacity relative that directly contributed to pathology.

The results from this study motivate future investigations. By combining the depth offered by bulk repertoire sequencing with the resolution of single-cell repertoire and gene expression, STAIR analysis allows for the characterization of B cells present at multiple time points in detail. Extending these observations to studies where anti-CD20 therapy has strong evidence for efficacy, such as MS may be particularly worthwhile. These studies could test the hypothesis that the depletion of common B cell subsets may predict efficacy across diseases. Applying STAIR analysis to the simultaneous investigation of other therapies, such as anti-CD19 therapy (inebilizumab)—recently shown to be efficacious for the treatment of NMO—could allow for the optimal selection of B cell depleting agents for preventing disease relapse (77). While RTX is a reliable treatment modality for MuSK MG, data for AChR MG has been mixed and patient relapses are common (12). An investigation of factors that contribute to relapse after RTX in the context of AChR MG using similar approaches could lead to the development of improved therapeutics for diagnosing and treating patients with MG regardless of disease subtype.

## Methods

### Patient-derived specimens

Peripheral blood was collected from MG subjects (Yale Myasthenia Gravis Clinic, New Haven, Connecticut, USA). A MuSK MG cohort (n = 3) was identified for purposes of investigation of post-RTX relapses with these characteristics that we have described previously: (i) a RTX-

induced CSR/MM (complete stable remission, minimal manifestation) clinical status (>1 year), (ii) repopulation of the B cell compartment after RTX, and (iii) disease relapse following sustained CSR/MM after rituximab (24, 25). Selected patients may have been treated with RTX prior to study entry. All pre-RTX sample collection time points precede post-RTX time points by at least one infusion of RTX. An additional asymptomatic (by Myasthenia Gravis Foundation of America, that is, MGFA classification score) AChR MG control subject was also included in our study.

### Bulk library Preparation

RNA was prepared from frozen peripheral blood mononuclear cells using the RNAEasy Mini kit (Qiagen) per manufacturer’s instructions. Cells were pelleted and lysed in freezing media without washing to preserve cell populations sensitive to cryopreservation. RNA libraries were made using reagents supplied by New England Biolabs as part of the NEBNext ® Immune Sequencing Kit. The RNA was reverse-transcribed into cDNA using a biotinylated oligo dT primer. An adaptor sequence, containing a universal priming site and a 17-nucleotide unique molecular identifier (UMI) was added to the 3’ end of all cDNA. Following purification using streptavidin-coated magnetic beads, PCR was performed to enrich for immunoglobulin sequences using a pool of primers targeting the IGHA, IGHD, IGHE, IGHG, and IGHM regions. This immunoglobin-specific primer pool contained tailed sequences with a priming site for a secondary PCR step. The second primer is specific to the adaptor sequence added during the RT step, and contains a sample index for downstream pooling of samples prior to sequencing.

Following purification of PCR products using AMPure XP beads (Beckman), the secondary PCR was performed in order to add the full-length Illumina P5 Adaptor sequence to the end of each immunoglobin amplicon. The number of secondary PCR cycles was tailored to each sample to avoid entering plateau phase, as judged by a prior quantitative PCR analysis. Final products were purified, quantified with a TapeStation (Agilent Genomics) and pooled in equimolar proportions, followed by high-throughput 2×300 base-pair paired-end sequencing with a 20% PhiX spike on the Illumina MiSeq platform according to manufacturer’s recommendations, except for performing 325 cycles for read 1 and 275 cycles for read 2.

### Raw read quality control and assembly for bulk libraries

Bulk repertoire data processing and analysis was carried out using tools in the Immcantation framework (http://immcantation.org). Preprocessing was carried out using pRESTO v0.5.10 (78) as follows. 1) Reads with a mean Phred quality score below 20 were removed. 2) Reads were aligned against constant region primer and template switch sequences, with a maximum mismatch rate of 0.2 and 0.5 respectively. Reads failing to match both a constant region primer and template switch sequence were removed. 3) Remaining reads were grouped based on unique molecular identifier (UMI) sequences determined by the 17 nucleotides preceding the template switch site. 4) Separate consensus sequences for the forward and reverse reads within each UMI group were constructed with a maximum error score of 0.1 and minimum constant region primer frequency of 0.6. If multiple constant region primers were associated with a particular UMI group, the majority primer was used. 5) Forward and reverse consensus sequence pairs were aligned to each other with a minimum overlap of 8 nucleotides and a maximum mismatch rate of 0.3, and assembled into full length sequences. Sequence pairs that failed to align to each other were assembled by alignment against the IMGT human germline IGHV reference database (IMGT/GENE-DB v3.1.19; retrieved June 21, 2018; Giudicelli et al. 2005) with a minimum overlap of 0.5 and a E-value threshold of 1×10-5. 6) Isotypes were assigned by local alignment of the 3’ end of each consensus sequence to isotype-specific constant region sequences with a maximum mismatch rate of 0.4. 8) Duplicate sequences were removed, except those spanning multiple biological samples and/or with different isotype assignments. Constant region primers, isotype-specific internal constant region sequences, and template switch sequences used are available at: https://bitbucket.org/kleinstein/immcantation/src/default/protocols/AbSeq.

### Single-cell library preparation and gene expression analysis

B cells were isolated from PBMCs by Human Pan B cell immunomagnetic negative selection kit (StemCell) per manufacturer’s instructions. Additional fluorescence-activated cell sorting (FACS) was performed for bead sorted B cells from MuSK MG post-RTX derived samples. B cells were labelled with 7AAD Viability Staining Solution (BioLegend) and 7AAD positive populations were removed. Additionally, B cells were labelled with FITC-labelled anti-IgD antibody (BioLegend clone IA6-2) per manufacturer’s instructions and FITC positive populations were removed. Of note, the purpose of this additional filtering was the removal of all naïve B cell populations. Sorted B cells were loaded into the Chromium Controller (10x Genomics) to form emulsion droplets. Libraries were prepared using the Chromium Single-cell 5′ Reagent Kit (10x Genomics) for version 3 chemistry per the manufacturer’s protocol. Samples were sequenced on the NovaSeq with 100×100 or 150×150 paired end reads for gene expression and BCR libraries respectively. Base calls were converted to fastq sequences and demultiplexed using the cellranger mkfastq function from 10X Genomics 2.2.0 and aligned to the GCRhg38 genome supplied by 10X Genomics. Sparse count matrices, barcode assignments and feature calls were made using the cellranger count function. The resulting output was loaded into Seurat v.2.3.4 for analysis. Cells with less than 50 genes detected, or mitochondrial content above 0.2 of all transcripts were discarded. Individual cells expressing detectable CST3, CD3E, KLRB1, NKG7 and GATA2 transcripts were excluded to remove non-B cell populations like T cells and myeloid cells. Single-cell expression values were computed from log normalized count values (79). 12,534 cells were identified in the final filtered analysis. Clusters were assigned and tSNE was computed after first scaling the data, removing cells with elevated mitochondrial content, identifying highly variable genes using FindVariableGenes and correcting for batch effects using the first 20 dimensions from canonical correlation analysis as identified by visualizing with MetageneBicorPlot. The first 10 dimensions were used for PCA based on the location of the inflection point from PCElbowPlot. FindClusters was then used to identify clusters in Seurat v.2 which uses shared nearest neighbor clustering after PCA (79).

Cell subsets were identified using a “basis set” of published marker genes called immunoStates (80). The mean expression of each gene was computed for each cluster from the scaled log normalized expression values; each cluster was then assigned to the immunoState subset with the highest Pearson correlation value. The absence of transitional B cells in the immunostates dataset meant this cluster was identified manually using marker genes alone. Small clusters with elevated expression of mitochondrial associated genes were also excluded from further analysis. We confirmed our assignments by visualizing the expression of known gene expression markers for instance, CD27 (for memory B cells), CD20 (for B cells in general except plasma cells), CD10 (for transitional B cells) and BLIMP1 (for plasma cells) (Figure S6A-C). Overall marker expression was also visualized based on the percent of cells in each cluster expressing the gene and the overall mean level of gene expression in each cluster (Figure S6D,E). To further confirm the identities of these single-cell clusters, we assessed their paired IGH repertoire features for mean somatic hypermutation (SHM) frequency and isotype usage which were consistent with expected values (Figure S6F,G).

### Bulk BCR repertoire V(D)J gene annotation, additional sequence filtering, clonal assignment and germline reconstruction for Single-cell tracing of adaptive immune repertoires (STAIR)

V(D)J germline assignments were made with IgBLAST v1.7.0 (81) using the June 21, 2018 IMGT reference gene database from bulk libraries. Reconstructed V(D)J sequences from single cell sequencing were extracted using the cellranger vdj function from fastq reads. V(D)J germline segments were re-assigned using IgBLAST v.1.7.0 also using the June 21, 2018 version of the IMGT gene database. Cells with multiple IGH V(D)J sequences were assigned to the most abundant IGH V(D)J sequence by UMI count. Single-cell V(D)J BCR repertoires as well as previously published single-cell PCR (scPCR) derived V(D)J sequences were pooled with bulk V(D)J BCR repertoires (24, 25). Following V(D)J annotation, non-functional sequences were removed from further analysis and functional V(D)J sequences were assigned into clonal groups using Change-O v0.3.4 (30). Sequences were first partitioned based on common IGHV gene annotations, IGHJ gene annotations, and junction lengths. Within these groups, sequences differing from one another by a length normalized Hamming distance of 0.126 were defined as clones by single-linkage clustering. This distance threshold was determined based on the average local minima between the two modes of the within-sample bimodal distance-to-nearest histogram for the three patients across multiple time points; thresholds were determined using a kernel density estimate in the same manner described previously (30). Germline sequences were then reconstructed for each clonal cluster (VH) with D segment and N/P regions masked (replaced with Ns) using the CreateGermlines.py function within Change-O v0.3.4.

### Differential gene expression analysis

Single-cell log normalized expression values were imputed using the ALRA algorithm to account for dropout during differential gene expression analysis (82). The mean value for each gene per sample was subtracted, the mean expression value for cells of a given status and sample was computed and values were z-score normalized for each patient. Student’s t-tests without pairing were performed to assign p-values for hypothesis testing. False detection rate (FDR) correction was performed using Storey’s method implemented within the qvalue package v3.9 in R from Bioconductor on the 12,066 p-values extracted from our analysis of cluster 3 associated genes (83). Gene ontologies were assigned using the enrichr package v1.0 in R which implements a wilcoxon signed rank test for significance (84). A q-value (corrected p-value) threshold of 0.05 was used for assigning significance for all tests of significance.

### Analysis of SHM, diversity and clonal overlap from BCR repertoires

Mutations relative to the germline sequence were quantified using ShazaM v0.1.8 in R v3.4.2 (85). Diversity analysis was performed using Alakazam v0.2.11 and assessed using the generalized Hill index, with uniform downsampling (to the number of V(D)J from the sample with the fewest sequences to account for different sequencing depth) and 2,000 replicates of bootstrapping from the inferred complete clonal abundance (86). Clonal overlap was computed using a Bray-Curtis metric implemented by the function scipy.spatial.distance.braycurtis in scipy v1.1.0 (87).

### Inference of B cell lineage trees and prediction of intermediate timepoints

B cell lineage tree topologies and branch lengths were estimated using the dnapars program distributed as part of PHYLIP (v3.697) (88). A maximum parsimony algorithm was used to label the internal nodes of each lineage tree as either or pre-RTX, post-RTX, given the date of sampling of each sequence at the tree’s tips. This was accomplished using an implementation of the Sankoff parsimony algorithm (89) in which switches from post-RTX to pre-RTX are forbidden using a cost matrix that weights post-RTX to pre-RTX switches as 1000 times more costly than switches in the opposite direction. Clusters of internal nodes separated by zero length branches (polytomies), were re-ordered using nearest-neighbor interchange (NNI) moves to minimize the number of label switches along the tree. These analyses were performed using IgPhyML v1.10 (90).

### Statistics

R v.3.4.2 (85) and Python 3.5.4 was used for all statistical analysis. Dataframe handling and plotting was performed using functions from the tidyverse 1.2.1 in R and pandas 0.24.2, scipy 1.1.0 and matplotlib 2.2.2 in python. All parametric statistical testing aside from that used for differential gene expression analysis (described previously) was performed in R using the aov function for two-way ANOVA or t.test functions for paired two-tailed student t-tests (or one-tailed student t-test when appropriate). A significance threshold of <0.05 was used and shown on plots with a single asterisk; double asterisks correspond to a p <0.01 and triple asterisks correspond to a p<0.001.

### Study Approval

This study was approved by Yale University’s Institutional Review Board. Informed consent was received from all participating patients prior to inclusion in this study.

## Author contributions

R.J, P.S., S.H.K, and K.C.O. designed the study. R.J. and P.S. performed all the experiments. R.J. and K.B.H. analyzed the data. R.J, M.L.F., K.B.H, R.J.N, P.S., S.H.K, and K.C.O. wrote the manuscript. R.J.N, S.H.K, and K.C.O. provided resources, reagents, and funding.

## Acknowledgments

We thank Jason Vander Heiden for assistance with developing methods for processing V(D)J sequences from 10X Genomics sequencing. We thank Karen Boss and Abeer Obaid for helpful feedback on the text. We thank Drs. Eileen Dimalanta and Chen Song from New England Biolabs for providing reagents used for the preparation of bulk BCR libraries.

