## Supplemental Materials for "Single-cell repertoire tracing identifies rituximab refractory B cells during myasthenia gravis relapses"

| Patient | Age | Sex | Collection | Event Type | Antibody Titer | MGC Score | MGFA Class | MGFA PIS | PI Status Change | Prednisone  Dose (mg) |
| --- | --- | --- | --- | --- | --- | --- | --- | --- | --- | --- |
| 1 37 F | | | 6 | Pre-RTX | 2560 |  | 0 |  | Worse  (W) | 0 |
|  |  |  | 2 | Pre-RTX | 2560 |  | IIa |  | Exacerbation  (E) | 0 |
|  |  |  | 7 | Pre-RTX |  | 14 | IIIb |  | Worse  (W) | 0 |
|  |  |  | 10 | Post-RTX | 5.7 | 6 | IIb |  |  | 0 |
| 2 63 F | | | 3 | Pre-RTX | 2560 |  | IIb |  | Exacerbation (E) | 0 |
|  |  |  | 8 | Pre-RTX | 2560 | 0 | 0 | CSR* | Unchanged (U) | 0 |
|  |  |  | 9 | Post-RTX | 1280 | 7 | IIb |  |  | 0 |
| 3 53 F | | | 1 | Pre-RTX | 160 | 3 | IIa |  |  | 50 |
|  |  |  | 5 | Pre-RTX | <10 |  | IIa |  | Improved  (I) | 30 |
|  |  |  | 12 | Pre-RTX | <10 | 6 | IIa |  | Improved  (I) | 30 |
|  |  |  | 4 | Post-RTX | 0.89 | 3 | IIa |  |  | 10 |

Table S1. Demographic characteristics of MuSK MG study subjects at each collection time point. MGC Score and MGFA Class were collected for each time point. “MGFA Class” refers to Myasthenia Gravis Foundation of America classification system for MG symptoms. “MGC Score” refers to the Myasthenia Gravis Composite score. “Age” refers to patient age at time of “Post-RTX” relapse.

*Complete Stable Remission

| Patient | Collection | Relapse | Reads | Unique IgM | Unique IgG | Unique IgA | Clones |
| --- | --- | --- | --- | --- | --- | --- | --- |
| 1 | 6 | Pre-RTX | 321622 | 19552 | 25596 | 26067 | 20084 |
|  | 2 | Pre-RTX | 339025 | 11995 | 40274 | 33127 | 15015 |
|  | 7 | Pre-RTX | 518947 | 29872 | 23544 | 26715 | 31820 |
|  | 10 | Post-RTX | 810822 | 30492 | 12343 | 14239 | 31902 |
| 2 | 3 | Pre-RTX | 391364 | 4274 | 971 | 2617 | 5290 |
|  | 8 | Pre-RTX | 108953 | 9477 | 1513 | 4154 | 10088 |
|  | 9 | Post-RTX | 460288 | 13823 | 2475 | 7788 | 14630 |
| 3 | 1 | Pre-RTX | 364381 | 752 | 5785 | 2392 | 2427 |
|  | 5 | Pre-RTX | 508120 | 29 | 669 | 1288 | 239 |
|  | 12 | Pre-RTX | 188934 | 154 | 2321 | 4016 | 860 |
|  | 4 | Post-RTX | 477305 | 7366 | 12952 | 10060 | 7999 |

Table S2. Counts of reconstructed V(D)J sequences by isotype and clones from sequencing of pre-RTX and post-RTX bulk IGH repertoires.

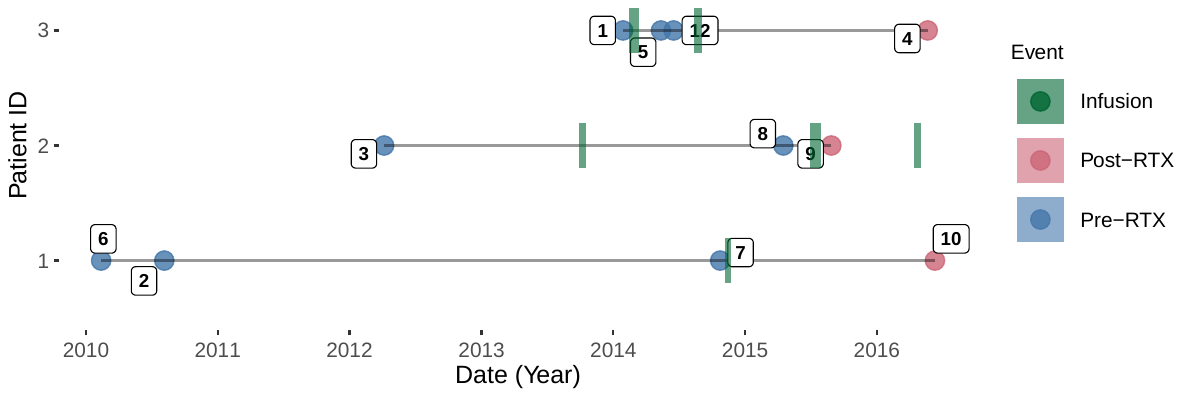

Figure S1. Timeline of three patients who experienced post-RTX relapses. Symptom score based on MGFA class and MGC scores were recorded at all collection time points (boxed labels).

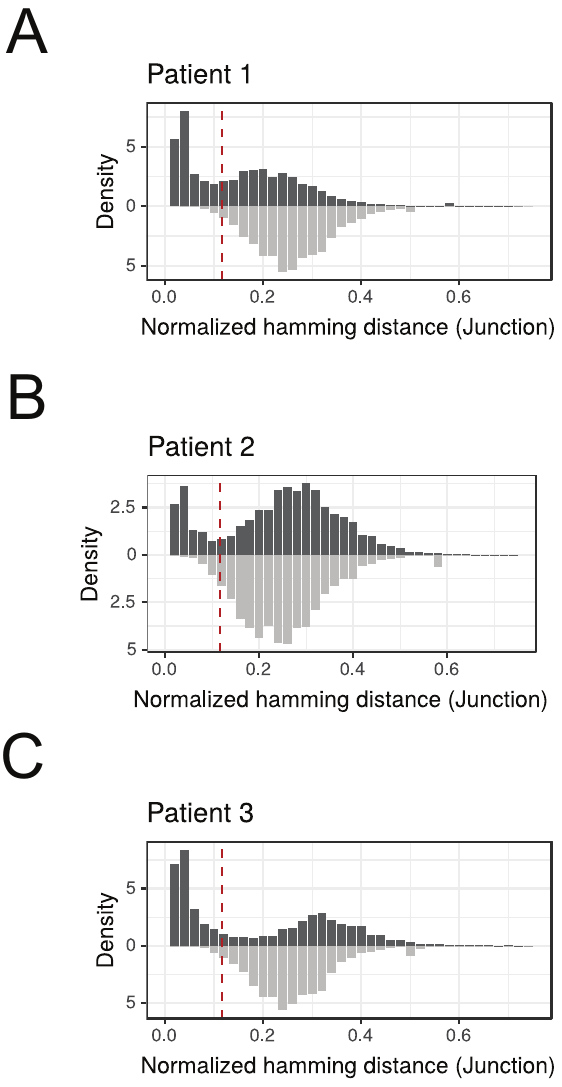

Figure S2. Distance-to-nearest plots used to identify a common threshold to use for hierarchical clustering based grouping of V(D)J sequences from high throughout sequencing of BCR repertoires. Red dashed lines corresponds to the threshold used for assigning clonal clusters.

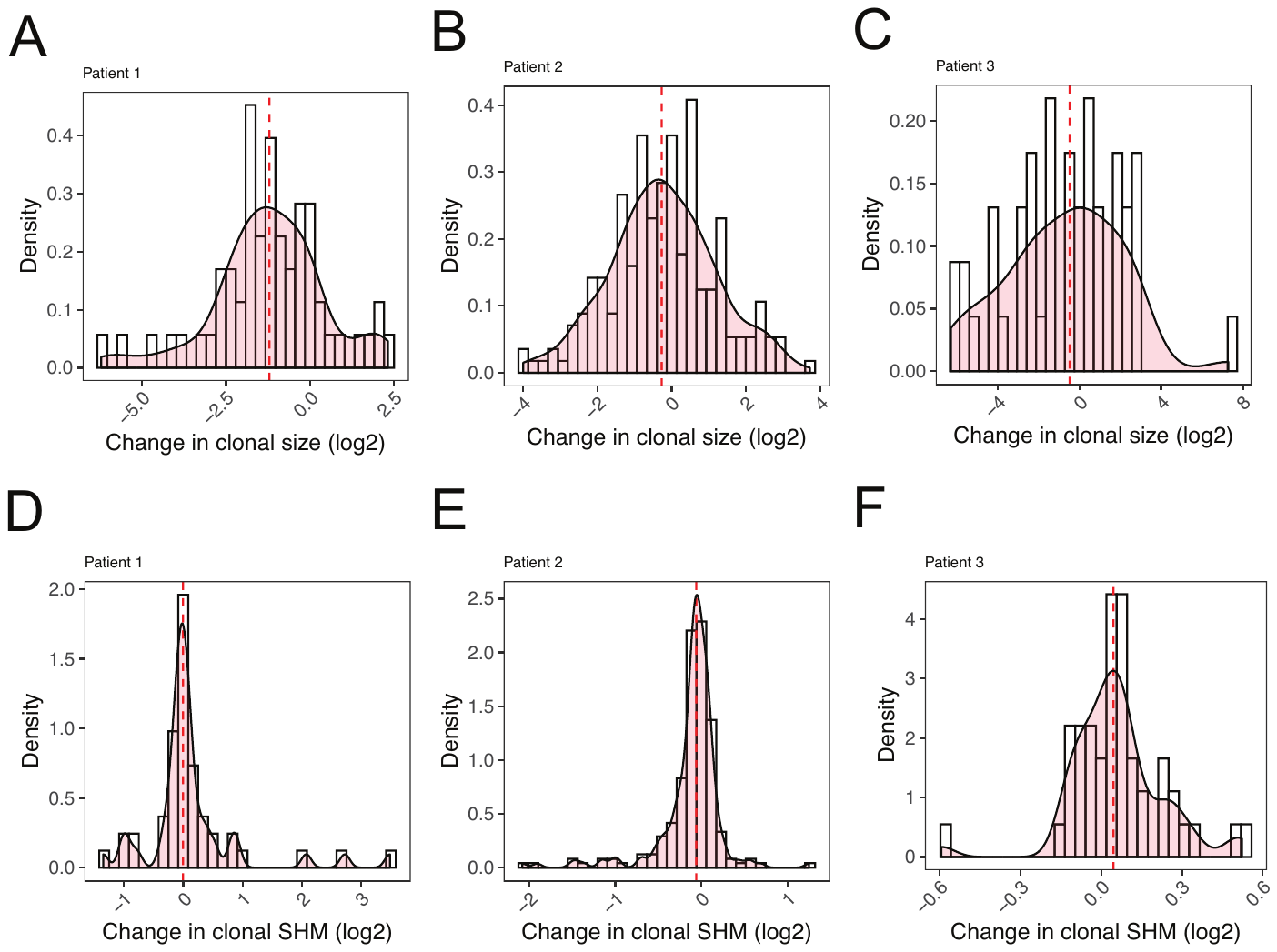

Figure S3. Persistient B cell clones do not show evidence of significant clonal expansion or accumulation of somatic hypermutations when comparing pre-RTX and post-RTX members. The presence of increases in overall somatic hypermutation for individual clones is quantified by comparing the average SHM frequency of each clone in the pre-RTX vs. post-RTX repertoire (A-C). Ratios are visualized as a histogram of the log2 fold changes in somatic hypermutation frequency values for these clones for each patient. Red dashed lines correspond to the median of each distribution. The presence of clonal expansions is evaluated by comparing the overall frequency of individual persistent clones at the post-RTX time point vs. the pre-rituximab time point (D-F). Frequencies are computed as a fraction of total unique V(D)J sequences and visualized in terms of a histogram of the log2 changes in total frequency for these clones. Red dashed lines correspond to the median of each distribution for each patient.

| Stathopoulos et al. 2017, BCR sequences from scPCR of circulating plasmablasts | | | |
| --- | --- | --- | --- |
| Patient | Clones | Traceable clones | Pre-RTX clones |
| 1 | 4 | 3 | 1 |
| 2 | 0 | 0 | 0 |
| 3 | 39 | 19 | 5* |
| Kazushiro et al. 2019, BCR sequences from scPCR of tetramer binding B cells | | | |
| Patient | Clones | Traceable clones | Pre-RTX clones |
| 1 | 22 | 3 | 1 |
| 2 | 4 | 0 | 0 |
| 3 | 12 | 2 | 0 |

Table S3. Count of plasmablast derived or tetramer-binding V(D)J sequences clonally related to members of the pre-RTX *or* post-RTX bulk BCR repertoire (Traceable clones) or to members of the pre-RTX *and* post-RTX bulk BCR repertoire (Pre-RTX clones). *One clone was observed to also correspond to a clone previously published to have specificity for MuSK autoantigen. The strongest binding member of this clone was annotated as “3-29” in the previous publication.

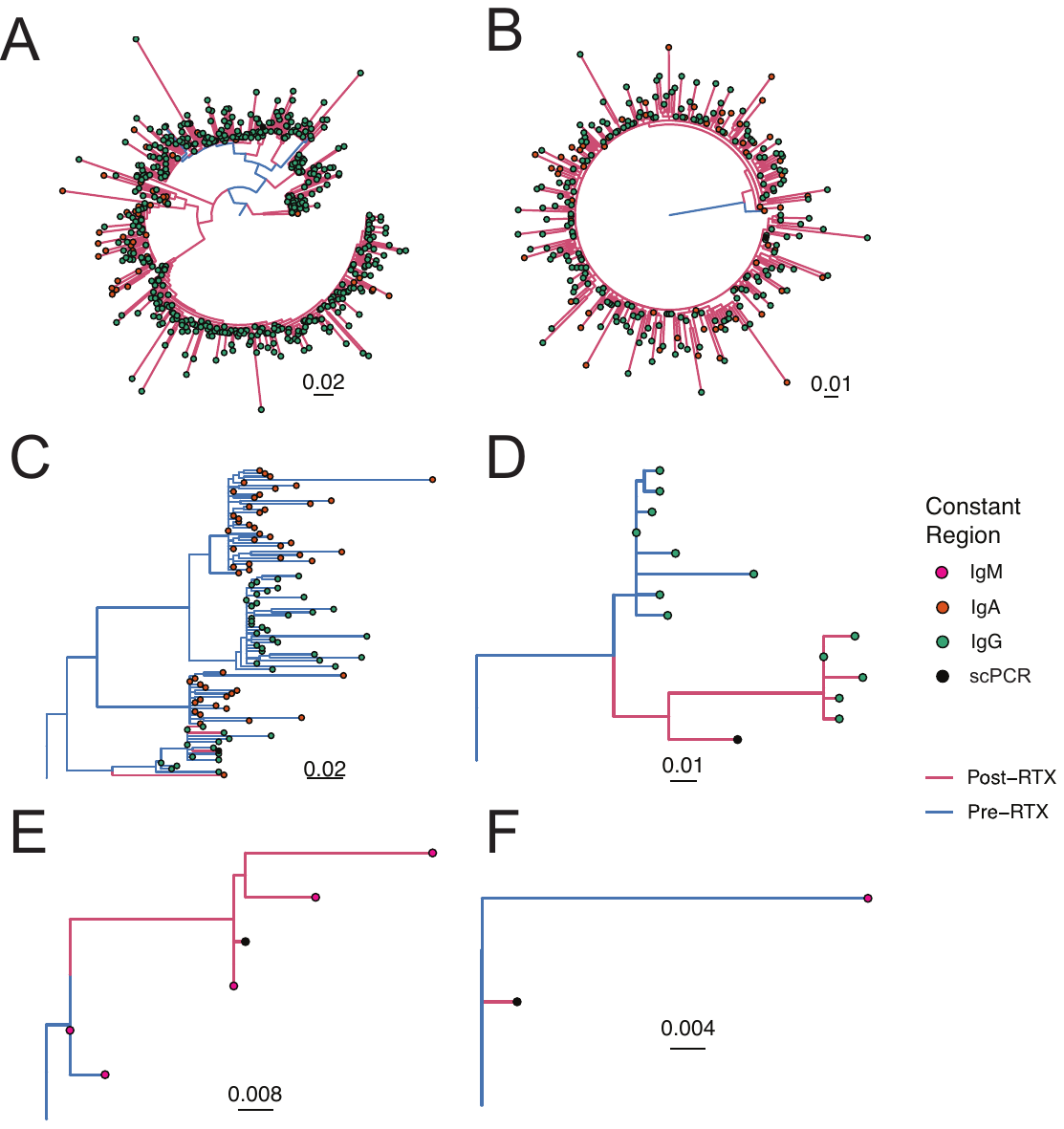

Figure S4. Clonal lineages containing scPCR-derived antibodies from Sanger sequencing identified in post-RTX repertoires. Maximum parsimony trees corresponding to clones that contain scPCR derived V(D)J sequences, and that have clonal variants present pre-RTX and post-RTX are presented (A-F). Sequences isolated using scPCR-based approaches from plasmablasts or by a fluorescent tetramer are denoted as "scPCR". Edge lengths are quantified based on intervening somatic hypermutations between observed V(D)J sequences per the scale. Colors correspond to whether each V(D)J sequence was collected from a pre-RTX or post-RTX time point (or both) and also the associated constant region.

| Patient | Collection | Status | Read Count | Cell Count | Mean Reads per Cell | Median Genes per Cell | VDJ Count |
| --- | --- | --- | --- | --- | --- | --- | --- |
| Control | MG295 | AChR MG Asymptomatic | 708747323 | 3580 | 197974 | 1499 | 3810 |
| 1 | MG189 | MuSK MG Post-RTX | 303494244 | 3550 | 85491 | 1189 | 2859 |
| 2 | MG139 | MuSK MG Post-RTX | 130365803 | 8502 | 15333 | 1130 | 6827 |
| 3 | MG192 | MuSK MG Post-RTX | 112260566 | 2501 | 44886 | 1790 | 1175 |

Table S4. Quality control from paired single-cell transcriptome and repertoire sequencing.

Figure S5. Example fluorescence activated cell sorting (FACS) gates for IgD^low^ B cells used as input for single-cell transcriptomics and BCR repertoire analysis. Example is shown for collection MG189.

| Cluster | Immunostates Assignment | Pearson Correlation | Final Assignment |
| --- | --- | --- | --- |
| 0 | naive_B_cell | 0.234 | Mature Naive |
| 1 | memory_B_cell | 0.125 | Memory |
| 2 | memory_B_cell | 0.122 | Memory |
| 3 | memory_B_cell | 0.166 | Memory |
| 4 | memory_B_cell | 0.251 | Memory |
| 5 | memory_B_cell | 0.351 | Memory |
| 6 | naive_B_cell | 0.136 | Transitional |
| 7 | not_assigned | -0.230 | Not Assigned |
| 8 | plasma_cell | 0.644 | ASC |
| 9 | memory_B_cell | 0.212 | Memory |
| 10 | not_b_cell | 0.063 | Not B cell |
| 11 | not_assigned | -0.135 | Not Assigned |

Table S5. Table of cluster assignments using the immunoStates basis set. Clusters were assigned to the B cell immunostate with the maximum pearson correlation coefficient when compared with the mean expression value of genes associated with each cluster. Unassigned clusters were manually annotated. One naive B cell cluster was assigned to a transitional B cell subset based on marker expression (cluster 6) and another cluster (cluster 10) was excluded owing to elevated expression of mitochondrial genes.

Figure S6. Assignment of B cell clusters based on single-cell transcriptome and repertoire according to known B cell subsets. Plotting of gene expression of key marker genes over t-SNE plot with intensity of shading correlated with amount of scaled log normalized expression for (A) CD27 (B) CD10 (MME) and (C) BLIMP1 (PRDM1). Dot plot of average log normalized expression (color) and fraction of cells expressing a set of marker genes (size) presented for B cell subsets derived from unbiased clustering (D) and when grouped into known B cell subset clusters (E). (F) Overall usage of different constant regions by B cell subset as a fraction of all V(D)J sequences associated with each B cell subset cluster per sample. Horizontal bars show the average frequency of constant region usage across patients. Frequencies belonging to the same patient are paired with a gray line. (G) Distribution of somatic hypermutation frequencies among all V(D)J sequences assigned to each B cell subset cluster. Horizontal bars show the average somatic hypermutation frequency for a given B cell subset cluster.

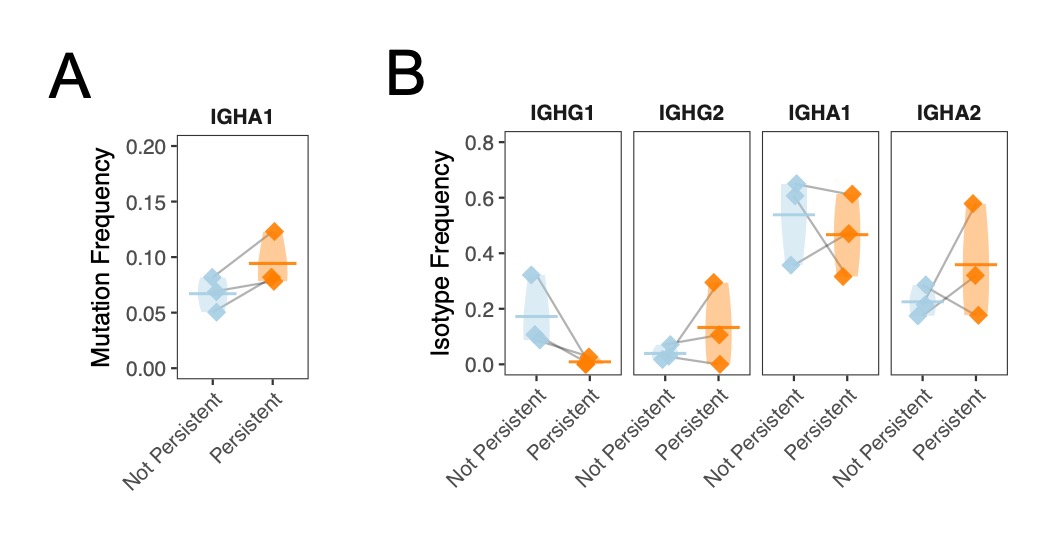

Figure S7. Persistent and non-persistent ASCs are associated with similar V(D)J repertoire. Overall constant region usage frequencies are quantified for persistent compared to non-persistent cells for the (A) ASC cluster per patient. Horizontal bars show the average frequency of constant region usage across patients. Frequencies belonging to the same patient are paired with a gray line. Individual SHM frequencies for persistent compared to non-persistent cells are presented for ASC cluster members (B). Only mean SHM frequencies are computed for isotypes with more than 3 V(D)J sequences. Horizontal bars show the average somatic hypermutation frequency for a given cluster. Frequencies belonging to the same patient are paired with a gray line.

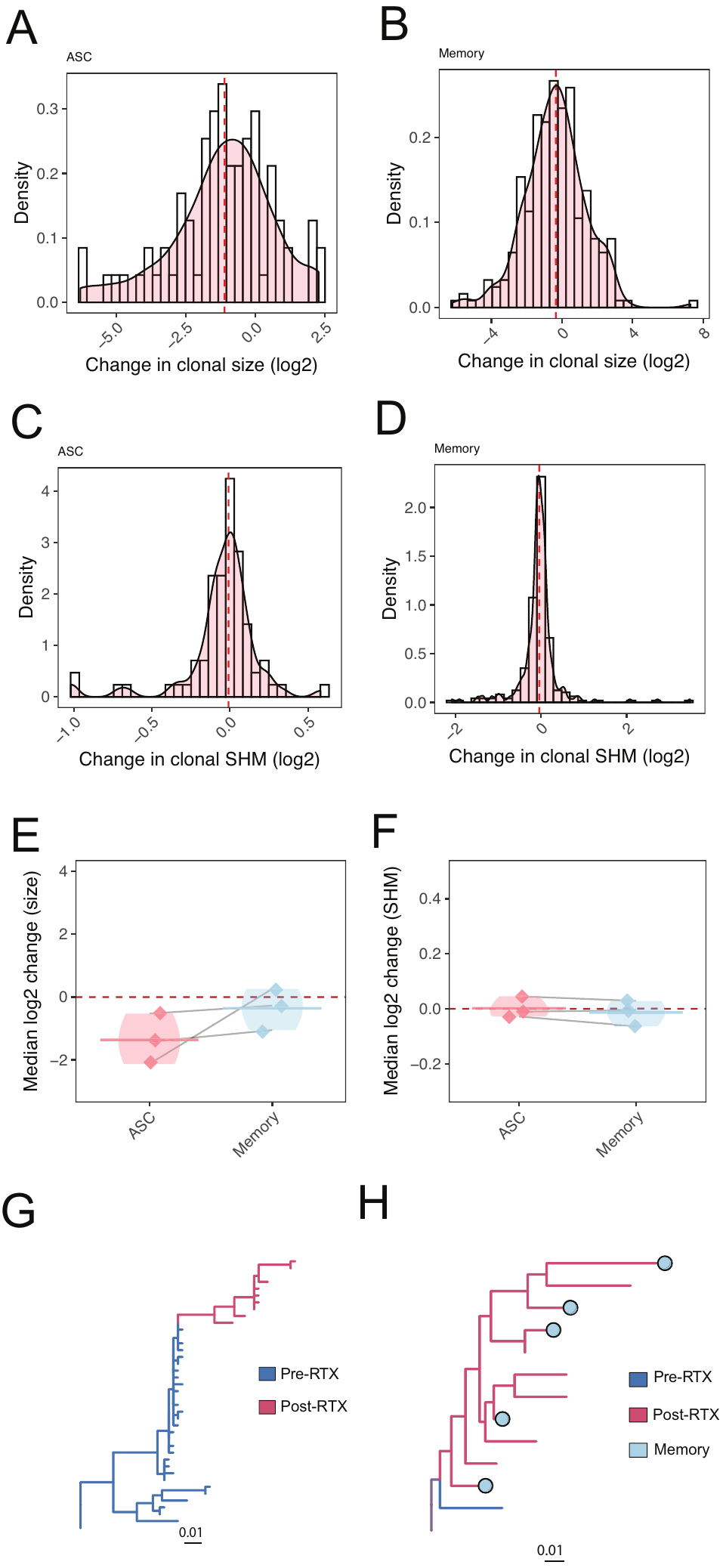

Figure S8. Persistent ASC and memory B cell clones do not show evidence of significant clonal expansion or accumulation of somatic hypermutations when comparing pre-RTX and post-RTX members. The presence of increases in overall SHM for individual clones is quantified by comparing the SHM frequencies averaged for each clone in the pre-RTX repertoire compared with post-RTX and visualized in terms of a histogram of the natural log change in somatic hypermutation frequency for each clone for each patient. Red dashed lines correspond to the median of each distribution for each patient. The set of clones examined is filtered for clones associated with ASCs (A) or memory B cells (B). The presence of clonal expansions is evaluated by comparing the overall frequency of individual persistent clones at the post-RTX time point compared to the pre-RTX time point from bulk IGH repertoires as a fraction of total unique V(D)J sequences and visualized in terms of a histogram of the natural log change in total frequency for these clones. Red dashed lines correspond to the median of each distribution for each patient. The set of clones examined is filtered for clones associated with ASCs (C) or memory B cells (D). The median of either the change (E) in clonal somatic hypermutation frequencies or clonal size distribution (F) associated with each patient is presented for each filtering strategy ("Total" for no filtering, "ASC" for filtering of ASC associated clones or "Memory" for filtering of Memory B cell associated clones. Horizontal bars show the median natural log change in clonal size or clonal somatic hypermutation frequency across patients. Values belonging to the same patient are paired with a gray line. Example B cell lineage trees consistent with continued somatic evolution after relapse are also shown; tree (G) was obtained from Patient 1, while tree (H) was obtained from Patient 2, and contains multiple memory B cells. Labels of internal nodes were predicted using the maximum parsimony algorithm described in Methods. The root of tree (H) is ambiguously Pre-RTX or post-RTX.

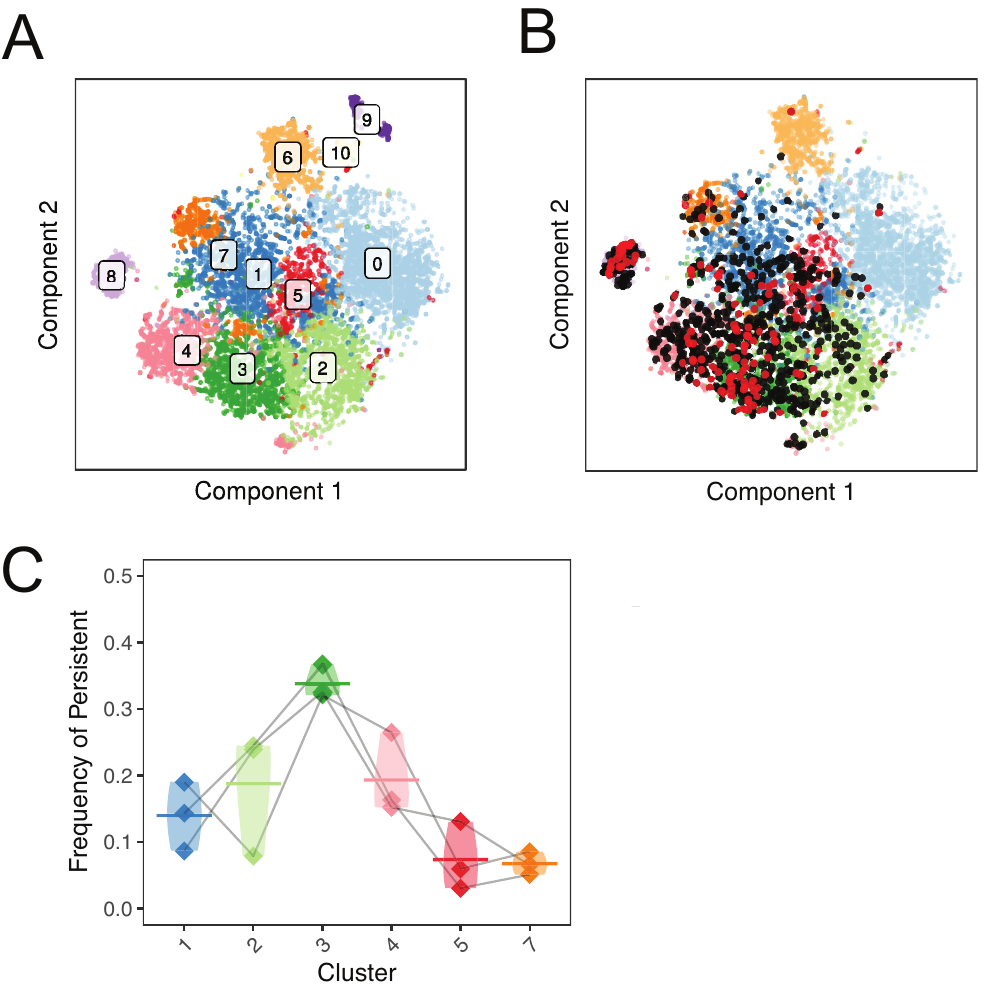

Figure S9. Identification of B cell clusters associated with B cell clonal persistence through unbiased analysis identifies subset associated with resistance to RTX depletion (cluster 3). Visualization of initial shared nearest-neighbor (SNN) clustering of circulating B cells by t-SNE with labels over clusters (A) or paired with B cell clones (black) or exact V(D)J sequence matches (red) (B) that match members of pre-RTX repertoires. (C) The frequency of each cluster among the set of persistent B cell clones is shown.

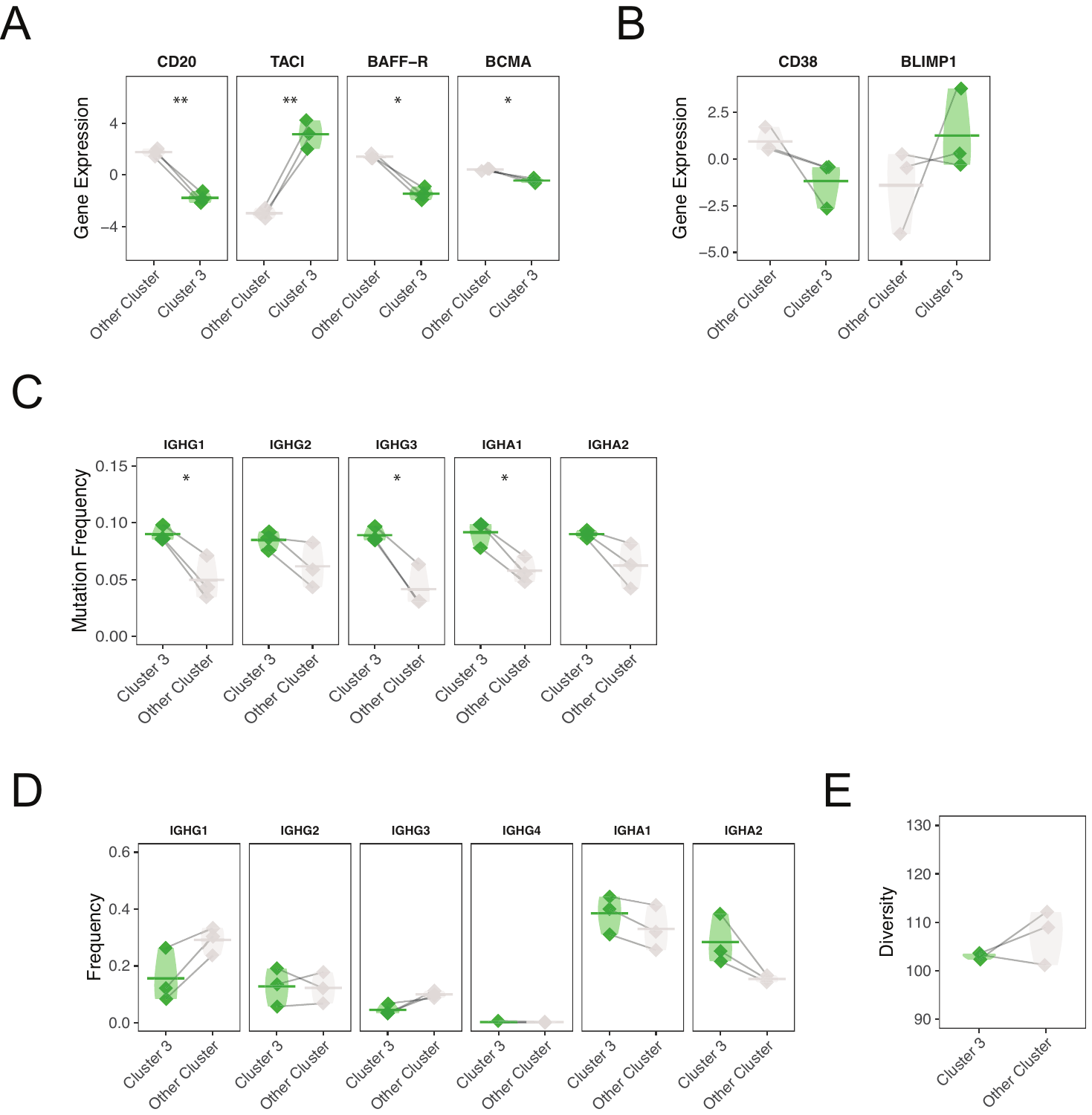

Figure S10. Cluster 3 has distinct transcriptional and V(D)J repertoire features. (A) The expression of the receptor target of rituximab CD20 (MS4A1) is presented for members of cluster 3 and members of other memory B cell clusters. The expression of known receptors for BAFF and APRIL (TACI or TNFRSF13B; BAFF-R or TNFRSF13C; BCMA or TNFRSF17) is also presented. (B) The expression of plasma cell associated genes (CD38; BLIMP1 or PRDM1) is also presented for members of cluster 3 and members of other memory B cell clusters. Normalized gene expression values are computed from scaled log normalized counts of barcoded transcripts aligned to each gene. Horizontal bars show the average normalized gene expression across patients. Expression values belonging to the same patient are paired with a gray line. (C) Mean somatic hypermutation frequencies for cluster 3 compared to other memory B cells is quantified per patient. Horizontal bars show the average somatic hypermutation frequency for a given set of cells. Frequencies belonging to the same patient are paired with a gray line. (D) Overall constant region usage frequencies are quantified for cluster 3 compared to other memory B cells per patient. Horizontal bars show the average frequency of constant region usage across patients. Frequencies belonging to the same patient are paired with a gray line. (E) Simpson’s diversity of cluster 3 members compared to other memory B cell cluster members.

Data file S1. List of differentially expressed genes for cluster 3 compared to all other memory B cell clusters. Only differentially expressed genes with an adjusted p-value less than 0.05 with FDR correction by Storey’s method are presented.

See attached cluster3.csv

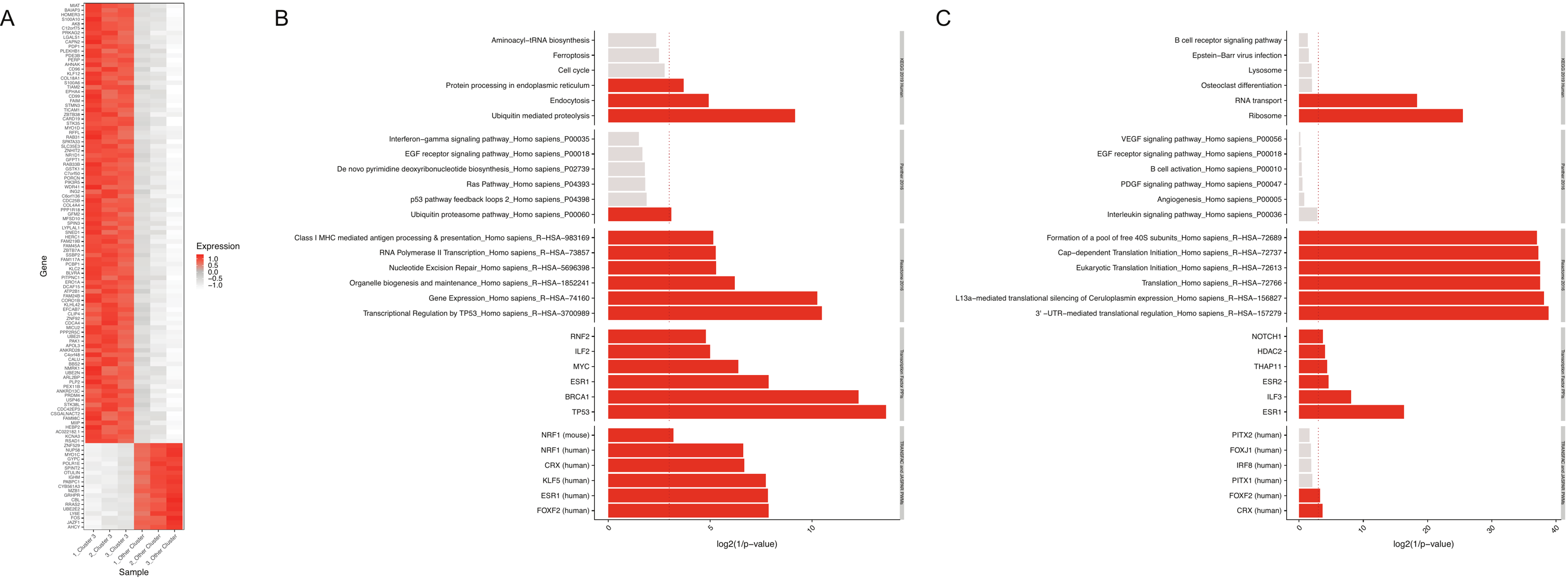
Figure S11. Cluster 3 memory B cells have a MYC and p53/BRCA1 signature. (A) Heatmap of significantly differentially expressed genes for cluster 3 compared to other memory B cells, with a greater than >0.4 z-score difference between persistent and non-persistent labels. enrichR gene ontology analysis of either genes (B) up-regulated or (C) down-regulated among cluster 3 memory B cells compared to other memory B cells. Only differentially expressed genes with an adjusted p-value less than 0.05 with FDR correction by Storey’s method were evaluated. Red bars correspond to significantly associated gene ontology assignments (p < 0.05 by wilcoxon signed rank test).
